# Multiomics reveal that silk fibroin and sericin differentially potentiate the paracrine functions of mesenchymal stem cells and enhance tissue regeneration

**DOI:** 10.1101/2022.09.29.510013

**Authors:** Yanan Zhang, Renwang Sheng, Jialin Chen, Hongmei Wang, Yue Zhu, Zhicheng Cao, Xinyi Zhao, Zhimei Wang, Chuanquan Liu, Zhixuan Chen, Po Zhang, Baian Kuang, Haotian Zheng, Qingqiang Yao, Wei Zhang

## Abstract

Silk fibroin (SF) and sericin (SS), the two major proteins of silk, are attractive biomaterials that show great potential in regenerative medicine. However, their biochemical interactions with stem cells were not fully understood. Here, we employed multiomics to obtain a global view of the triggered cellular processes and pathways of MSCs by SF and SS. Integrated RNA-seq and proteomics revealed that SF and SS strongly enhanced the paracrine activity of MSCs through differentially activating integrin and glycolytic pathways, rather than directly regulating stem cell fate to initiate multiple but distinct biological processes in MSCs. Those specific paracrine signals of MSCs stimulated by SF and SS effectively promoted skin wound healing by influencing the behaviors of multiple resident cells in skin wound microenvironments. This study provides comprehensive and reliable insights into the cellular interactions with SF and SS, enabling future development of silk-based therapeutics for tissue engineering and stem cell therapy.

## 1. Introduction

The shortage of tissues/organs for transplantation has always been a major concern worldwide (*1*). Tissue engineering is emerging as a potential alternative to these problems, which involves the effective integration of cells, scaffolds, and signals, referred to as a tissue engineering triad, for the development of biological substitutes to restore, maintain and replace the injured tissues/organs (*2*). Scaffold plays a significant role in the field of tissue engineering, which provides structure and substrate for cells to attach and growth, and subsequent tissue formation (*3*). A variety of natural or synthetic biomaterials have been explored to develop tissue engineered scaffolds. By incorporating exogenous growth-stimulating signals such as growth factors or small molecules, the scaffolds can provide specific bioactivities, i.e., biochemical signals required for cellular behaviors and tissue regeneration (*4*). As biomaterials are essentially comprised of chemical molecules, they are indubitably capable of supplying inherent biochemical signals to guide and influence the cells. However, the intrinsic biochemical signals of biomaterials are usually ignored and their global effects on the modulation of cellular functions and tissue regeneration, remain to be elucidated, particularly at an in-depth molecular level (*5*).

Silk derived from silkworm *Bombyx mori* is a natural protein polymer, which has been used clinically as surgical sutures for centuries (*6*). In the past decades, silk has attracted widespread attention in the fields of tissue engineering and regenerative medicine, due to its good biocompatibility, remarkable physical properties, and non-cytotoxic degradation products (*7*). Chemically, silk is mainly made of two kinds of proteins, an outer coating of silk sericin (SS) and a central core of silk fibroin (SF), which account for about 25–30% and 70–75% of total silk weight, respectively (*8*). SF is extracted from the raw silk by complete removal of SS, a process called degumming. The degummed silk (i.e., SF) has long been recognized as a promising biomaterial and has been widely used for tissue engineering applications. We and other researchers have developed various formats of SF scaffolds for the repair and regeneration of bone, cartilage, skin, cornea, etc (*9–12*). In contrast, SS was previously regarded as a waste material during degumming and was found to cause immunological responses when combined with SF (*13*). However, neat SS was later found as an emerging biomaterial with good biocompatibility and low immunogenicity (*14*). Specifically, many studies have reported the successful application of SS-based scaffolds for bone, skin, cartilage, and myocardial tissue regeneration (*15, 16*). Thus, both SF and SS have extensively been exploited as potential biomaterials in tissue engineering and regenerative medicine applications.

In particular, early studies reported that SF and SS biomaterials can provide biochemical signals to exert a multitude of bioactivities to stimulate cell migration, adhesion, proliferation, and even differentiation of various cell types (*16–18*). Martínez-Mora et al. reported that both SF and SS supported the migration of MDA-MB-231 and Mv1Lu cells by activating the MEK, JNK, and PI3K signaling pathways (*18*). Meng et al. found that the ratio of SF and SS in electrospun films regulated the macrophage polarization towards M1/M2 phenotypes (*19*). Notably, some studies have shown that SF-based scaffolds could regulate the proliferation and multi-lineage differentiation (e.g., osteogenic, chondrogenic, and endothelial differentiation) of stem cells, while other studies reported inconsistent results that SF lacks the bioactivity to guide stem cell differentiation (*20–23*). On the other hand, it has widely been reported that SS improved cell proliferation and maintained the stemness of stem cells, thereby being used for serum-free cell culture (*16*). However, recent studies also found that SS-incorporated scaffolds could induce the directional differentiation of stem cells, such as osteogenic and epithelial differentiation (*24, 25*). In general, both SF and SS exhibit multiple bioactivities that can regulate cellular behaviors and facilitate the repair and regeneration of various tissues, while their effects seemed to be multifaceted and controversial, especially in stem cell differentiation. Besides, how SF and SS interact with cells, especially stem cells, what are the dominant cellular processes initiated by SF and SS, and which one of SF and SS is more suitable for the repair and regeneration of specific tissues, are still not clear and need to be further explored. Therefore, more in-depth studies with advanced techniques should be performed to identify the global cellular responses to SF and SS, which will help predict their most possible biological effects in the body and thereby precisely and effectively guide tissue regeneration.

Previous studies have used molecular biology techniques such as qPCR and microarrays to evaluate the effects of SF and SS on cells, but these techniques are limited by low throughput and reproducibility (*15, 17, 26*). In recent years, the emergence and development of “omics” techniques has overcome the limitation of traditional molecular biology techniques and allowed us to understand the complicated cell-biomaterials interactions (*27, 28*). Specifically, transcriptomics and proteomics are capable of providing an opportunity to unbiasedly view the global cellular activities to SF and SS and obtain critical insights on the influenced biological processes and cellular pathways (*29*). It is worth mentioning that transcriptomics often reflects more global but less reliable cellular changes at the gene expression level, whereas proteomics can provide more reliable but less global cellular responses at the protein expression level, thus the integrated transcriptomics and proteomics could complement each other to obtain comprehensive and reliable insights into the biochemical interactions and processes mediated by SF and SS (*30, 31*). Recent studies have used RNA sequencing (RNA-seq) or proteomic analysis to investigate the cell-biomaterials interactions and revealed the regulation of stem cell behaviors by nanoclay, molybdenum disulfide, or bioceramics (*27, 32–35*). However, no studies have explored the cellular interactions with SF or SS at the whole transcriptome or protein levels.

In the present study, we employed integrated multiomics to comprehensively investigate the biochemical interactions of SF and SS with mesenchymal stem cells (MSCs) to elucidate the triggered cellular responses and pathways. The integrated transcriptomics and proteomics revealed multiple biological processes of MSCs with widespread gene and protein changes initiated by SF and SS. Specifically, both SF and SS strongly potentiated the paracrine activities of MSCs through differentially activating integrin and glycolysis signaling pathways, rather than directly regulating stem cell fate. Additionally, we investigated whether the paracrine functions of MSCs in response to SF and SS could mediate the local signaling processes in vivo and have implications for tissue regeneration. We found that SF and SS triggered specific paracrine signals of MSCs that promoted cell proliferation, migration, angiogenesis, immunomodulation, and skin wound healing processes. Moreover, the high consistency of transcriptomics and proteomics analyses in vitro and in vivo indicated the strong cellular responses of MSCs to SF and SS, as well as the stable and powerful biological effects of these paracrine signals. This work thus provides comprehensive and reliable insights into the cellular interactions with SF and SS via multiomics profiling, which enables future development of silk-based therapeutics for tissue engineering and stem cell therapy.

## 2. Results and Discussion

### 2.1 Preparation of SF and SS and their influence on the cellular behaviors of MSCs

In this study, both SF and SS were extracted from raw silk fibers, and the schematic of the extraction procedure was shown in Fig. 1A. The extracted SF and SS was yellowish and relatively clear with a concentration of ∼5% (w/v). Consistent with the previous report, SF and SS showed similar absorption peaks of amide I (SF: 1640.31 cm^-1^; SS: 1643.92 cm^-1^), amide II (SF: 1507.98 cm^-1^; SS: 1520.92 cm^-1^), and amide III (SF: 1231.04 cm^-1^; SS: 1245.68 cm^-1^) (Fig. 1B) (*36*). Moreover, SS (3277.43 cm^-1^) has a broader amide A band than SF (3275.02 cm^-1^), indicating the more random structure of SS and the higher crystallinity of SF (Fig. 1B) (*37*), which also reflected their distinct functions in natural silk fiber. These results demonstrated a successful preparation of SF and SS in our study.

**Figure 1.**
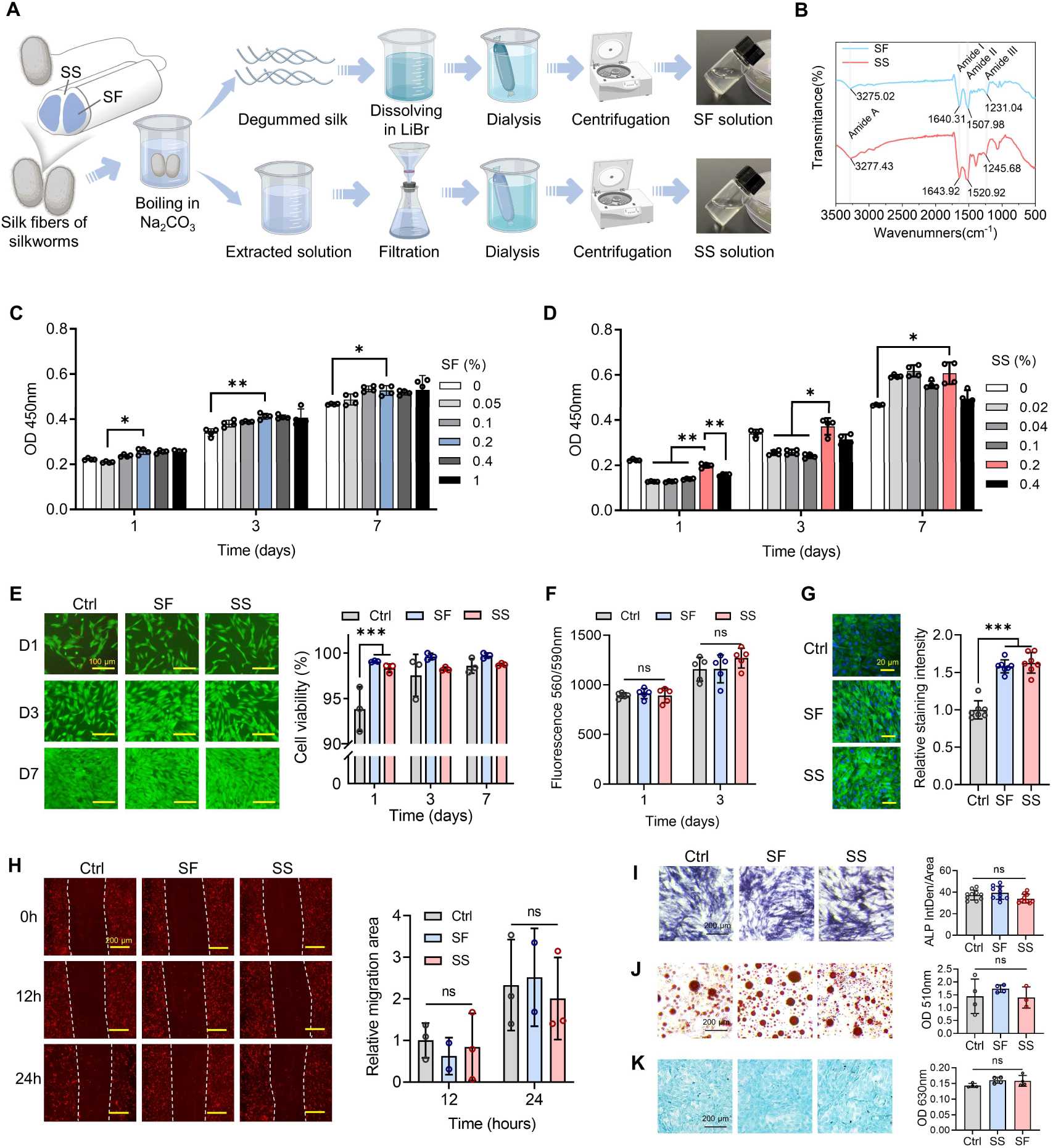
SF and SS regulate cellular behaviors of MSCs in vitro. (A) Schematic of the SF and SS extraction procedure. (B) FTIR spectrum of SF and SS. (C) Cell proliferation of MSCs treated with 0% to 1% (w/v) SF measured by CCK-8. (D) Cell proliferation of MSCs treated with 0% to 0.4% (w/v) SS measured by CCK-8. (E) Representative images and quantitative analysis of live/dead staining of MSCs treated with/without SF or SS. Calcium-AM for live cells (green) and PI for dead cells (red). Scale bars = 100 μm. (F) Alamar blue assay of MSCs treated with/without SF or SS. (G) Representative images and quantitative analysis of F-actin staining of MSCs treated with/without SF or SS. Scale bars = 20 μm. (H) Cell migration of MSCs treated with/without SF or SS using scratch assay. MSCs were pre-stained using Dil. Scale bars = 200 μm. (I-K) Representative images and quantitative analysis of ALP, oil red O and alcian blue staining of MSCs treated with/without SF or SS. Scale bars = 200 μm. The results were presented as mean ± SD. *P<0.05, **P<0.01, ***P<0.001.

MSCs are one of the most extensively investigated cell types in tissue engineering and stem cell therapy, with the capacity to differentiate into multiple lineages under different stimuli and culture conditions. Firstly, we investigated the influence of SF and SS on cellular behaviors of MSCs, including cell proliferation, viability, metabolic activity, morphology, migration, and differentiation. CCK-8 assay was conducted to evaluate the proliferation of MSCs when exposed to SF and SS with different concentrations. When cultured with SF (0.05% to 1%) or SS (0.02% to 0.4%) for 1, 3, and 7 days, MSCs in all groups could proliferate normally with time (Fig. 1C-D, S1). When exposed to 0.2% (w/v) SF, MSCs showed a slightly but significantly increased proliferation rate compared with the 0.05% (w/v) SF group on day 1 and the control group on days 3 and 7 (Fig. 1C). On days 1 and 3, the 0.2% (w/v) SS group had a significantly increased proliferation of MSCs compared to the SS groups with lower concentrations (0.002% to 0.1%) (Fig. 1D). Moreover, MSCs treated with 0.2% (w/v) SS exhibited a significantly enhanced proliferation compared with the control group on day 7 (Fig. 1D). Collectively, these results indicated that both 0.2% (w/v) SF and SS effectively promoted the proliferation of MSCs, which is consistent with the previous reports that SF and SS could act as bioactive molecules to support cell proliferation in tissue engineering applications (*38, 39*). Due to the higher proliferation levels of MSCs with 0.2% (w/v) SF or SS treatment, this specific concentration was chosen for the following experiments in this study.

Next, live/dead staining was performed to examine the cell viability of MSCs treated with SF and SS. After being cultured with/without SF or SS for 1, 3, and 7 days, MSCs in each group presented high cell viability (>93%) (Fig. 1E). Besides, it was observed that SF and SS significantly increased cell viability of MSCs on day 1 as compared to the untreated control, indicating the positive role of SF and SS in enhancing cell viability (Fig. 1E). However, this effect was not significant on days 3 and 7 when cell viability of each group was higher than 95%. Alamar blue assay was used to evaluate the effects of SF and SS on metabolic (mitochondrial) activities of MSCs, and no significant differences were identified between these three groups (Fig. 1F). We subsequently conducted F-actin staining to investigate the influence of SF and SS on the morphology of MSCs. The results indicated that MSCs in Ctrl, SF, and SS groups exhibited similar spindle-shaped morphologies (Fig. 1G). Although previous studies have reported that SF-based materials influenced cell morphology, it was primarily mediated by their physical properties, such as topography and stiffness, rather than their biochemical signals (*40, 41*). Interestingly, we found that MSCs treated with SF and SS displayed notably increased staining intensity of F-actin, indicating that SF and SS could promote F-actin formation in MSCs (Fig. 1G), which have been reported to improve cell adhesion and migration (*42, 43*). Therefore, we further investigated the migration of MSCs after SF and SS treatments via a scratch assay. However, the results indicated that the migratory capacity of MSCs was not significantly improved in response to SF or SS (Fig. 1H).

Importantly, MSCs are multipotent stem cells capable of differentiating into multiple lineages, such as osteogenic, adipogenic and chondrogenic lineages, playing a crucial role in tissue repair and regeneration. Therefore, we further evaluated the effects of SF and SS on the regulation of multi-lineage differentiation of MSCs. ALP, oil red O and alcian blue staining were conducted to evaluate the osteogenic, adipogenic and chondrogenic differentiation of MSCs treated with/without SF or SS, respectively. Unfortunately, no significant differences in ALP activity, lipid droplet formation or cartilaginous GAG production were detected between these three groups (Fig. 1I-K), suggesting that SF and SS treatments were not effective in directing cell differentiation of MSCs. Indeed, SF and SS are commonly considered as “bioinert’’ materials to actively regulate cell differentiation (*23, 44*). SF possesses only a few signaling peptide domains on its primary sequence, and SS is widely applied to serum-free medium to maintain cell proliferation and stemness of stem cells, suggesting that both SF and SS lack bioactive components to guide the directional differentiation of stem cells (*23, 44*). Although some previous studies have also reported that the composite scaffolds containing SF or SS were able to induce MSCs differentiation towards multiple lineages, including bone, cartilage, muscle, fat, and epithelium (*23, 24, 45-47*). However, the bioactivities of these scaffolds on cell differentiation were mainly attributed to the incorporation of exogenous bioactive molecules or the special physical properties (e.g., roughness, stiffness, and topography, etc), rather than the SF or SS biomaterials themselves.

Taken together, although multiple bioactivities of SF and SS have been reported before, our findings revealed that SF and SS only slightly enhanced the proliferation and viability of MSCs, while their effects on the metabolic activity, migration, morphology, and multi-lineage differentiation of MSCs seemed to be less significant. Moreover, the limited bioactivities of SF and SS in enhancing stem cell proliferation and viability seem insufficient to elucidate the compelling therapeutic effects of SF and SS in silk-based tissue engineering and regenerative medicine applications. Therefore, SF and SS probably influence other cellular functions of MSCs (e.g., paracrine signals), which needed to be further explored.

### 2.2 Widespread transcriptomic alterations triggered by SF and SS

RNA-seq is an emerging technology and has been widely used to determine widespread transcriptomic changes of stem cells in response to external stimuli, including chemical stimulus (e.g., agents and cytokines) and physical stimulus (e.g., radiation, stiffness, and topography) (*29, 32, 48, 49*). Therefore, RNA-seq was utilized to further investigate the global effects of SF and SS on the cellular responses of MSCs, which was helpful to identify their dominant bioactivities for tissue repair and regeneration.

#### 2.2.1 Global transcriptomic profile of MSCs treated with SF and SS

After being cultured with/without 0.2% (w/v) SF or SS for 3 days, RNA-seq was performed to investigate the widespread changes of MSCs in transcriptional level (Fig. 2A). Three replicates of Ctrl, SF, and SS groups were sequenced, and the gene expression levels were normalized by calculating the fragments per kilobase of transcript per million reads. The replicates of each group exhibited a high consistency (0.987 < r < 1 for Ctrl; 0.989 < r < 1 for SF; 0.985 < r < 1 for SS) (Fig. 2B). We performed a pairwise comparison between Ctrl, SF, and SS groups to identify differentially expressed genes (DEGs) of MSCs followed SF or SS treatment (Fig. 2C-F). The heatmap of all DEGs after clustering analysis revealed notable changes in the transcriptomic profile of MSCs between three groups (Fig. 2C). Specifically, 540 up-regulated DEGs and 1184 down-regulated DEGs were detected in the SF group compared to the Ctrl group (Fig. 2D). In the comparison of SS vs Ctrl, 487 genes were significantly up-regulated while 1675 genes were significantly down-regulated (Fig. 2E). These results indicated strong cellular responses of MSCs triggered by both SF and SS treatments. Moreover, 1411 up-regulated DEGs and 695 down-regulated DEGs were also detected between SF and SS groups, indicating their distinct influences on MSCs (Fig. 2F). The overlap of DEGs in paired comparisons showed that only 120 DEGs were identical while more than 1000 DEGs were especially changed in SF vs Ctrl (1111), SS vs Ctrl (1062), and SF vs SS (1160), confirming the robust but distinct effects of SF and SS on MSCs (Fig. 2G). Therefore, our results strongly suggested that SF and SS treatments led to remarkable cellular responses of MSCs, and their underlying mechanisms could be different depending on the DEGs.

**Figure 2.**
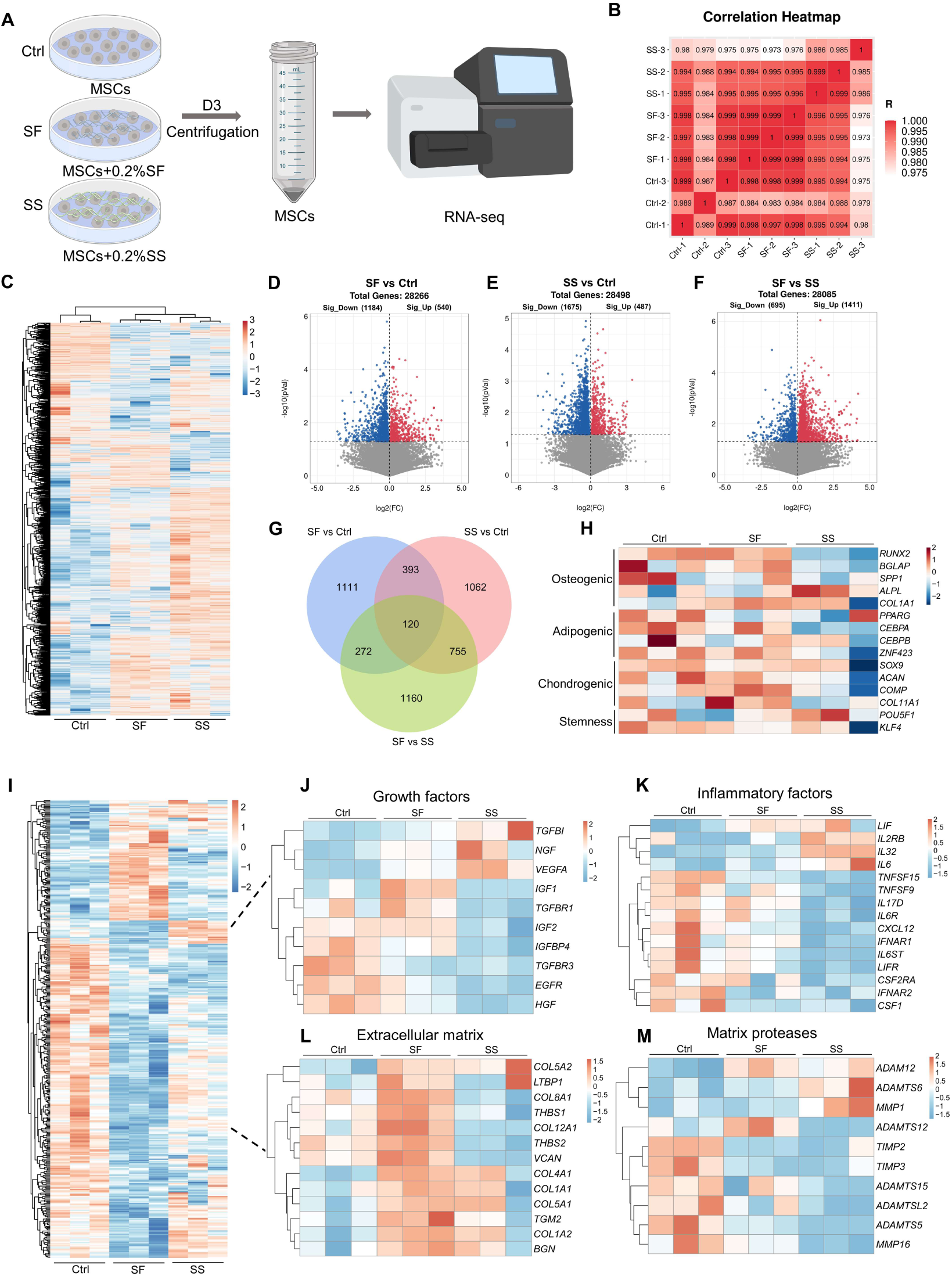
RNA-seq reveals widespread transcriptomic changes of MSCs triggered by SF and SS. (A) Schematic of the transcriptomic profiling of MSCs treated with/without SF or SS. (B) Correlation heatmap showed the relationship among the variables of Ctrl, SF, and SS groups. The correlation coefficient (R=0-1) represents the cluster analysis (P<0.05). (D-F) Volcano plots of gene expression levels in SF vs Ctrl, SS vs Ctrl, and SF vs SS groups. (G) Venn plot of identified DEGs in pairwise comparison. (H) Heatmap of gene expression of multi-lineage differentiation markers and stemness markers. (I) Heatmap of expressed genes encoding secreted proteins. (J-M) Heatmap of gene expression associated with growth factors (J), inflammatory factors (K), extracellular matrix (L), and matrix proteases (M).

Due to the attractive multi-lineage differentiation capacity of MSCs for tissue repair and regeneration, we firstly analyzed the gene expression levels of multi-lineage differentiation markers and stemness markers of MSCs after SF or SS treatment. It was observed that there was no significant difference in the expression of these marker genes related to osteogenic differentiation (*RUNX2*, *BGLAP*, *SPP1, ALPL*, and *COL1A1*), adipogenic differentiation (*PPARG*, *CEBPA*, *CEBPB*, and *ZNF423*), chondrogenic differentiation (*SOX9*, *ACAN*, *COMP*, and *COL11A1*), and stemness maintenance (*POU5F1* and *KLF4)* (Fig. 2H), confirming the “bioinert’’ property of SF and SS to guide stem cell differentiation (Fig. 1I-K).

In addition to the multi-lineage differentiation potential, the paracrine functions of MSCs, i.e., the capacity of secreting various bioactive molecules, including cytokines, growth factors, enzymes, and ECM components, has recently attracted considerable attention in the field of tissue engineering (*50–52*). Many studies have demonstrated that these paracrine products of MSCs could regulate various cellular behaviors of resident cells in local environments via paracrine signaling, and thereby play a critical role in tissue repair and regeneration (*50–52*). Moreover, increasing evidence indicated that the therapeutic effects of MSC transplantation were largely attributed to the paracrine functions of MSCs rather than their differentiation into tissue-specific cells (*51, 53*). Thus, the expression levels of genes encoding secreted proteins were evaluated in the identified genes from RNA-seq. A list of 3971 genes that encoded secreted proteins was obtained from THE HUMAN PROTEIN ATLAS (https://www.proteinatlas.org/), and 397 of them are differentially expressed between three groups, indicating that SF and SS had a strong influence on the secretome of MSCs (Fig. 2I). Specifically, the notable differences between Ctrl, SF, and SS were observed associated with growth factors, inflammatory factors, extracellular matrix (ECM), and matrix proteases (Fig. 2J-M), which were involved in the regulation of various biological processes for tissue repair and regeneration, including cell recruitment, stem/progenitor cell differentiation, angiogenesis, immunomodulation, and tissue remodeling (*50–52*). Collectively, SF and SS were more inclined to regulate the paracrine signals instead of the multi-lineage differentiation of MSCs, which could largely contribute to the therapeutic benefits of SF and SS in tissue regeneration.

In summary, both SF and SS triggered robust transcriptomic changes of MSCs, and the cellular changes induced by them were highly associated with the secretome of MSCs, which required further investigation to obtain the functional insights of these changes induced by SF and SS.

#### 2.2.2 Functional insights of cellular changes of MSCs treated with SF

To further explore the effects of SF on MSCs, Gene Ontology (GO) enrichment analysis for up-regulated DEGs (SF vs Ctrl) was performed using the DAVID database. Fig. 3A and S2A-B exhibited the top GO terms for biological processes (BP), cellular components (CC), and molecular function (MF) based on the P value, which revealed the notable effects of SF on MSCs. Several key GO terms were significantly enriched in BP, such as “extracellular matrix organization”, “collagen catabolic process”, “amino acid transport”, “collagen fibril organization”, “cellular response to amino acid stimulus”, “angiogenesis”, “leukocyte migration”, “cell adhesion”, “cellular response to hypoxia”, and “positive regulation of phosphatidylinositol 3-kinase (PI3K) signaling” (Fig. 3A). These results indicated that SF could regulate multifaceted functions of MSCs, including ECM deposition, collagen synthesis and secretion, angiogenesis, immunomodulation, and hypoxic adaptation (Fig. 3A). Subsequently, REVIGO analysis was conducted to refine the extensive significant GO terms of BP via reducing the redundancies of these terms and clustering the notable terms based on similarity measures (*32*). REVIGO analysis refined all enriched BP GO terms to angiogenesis, ECM organization, collagen catabolic process, cellular response to amino acid stimulus, amino acid transport, positive regulation of gene expression, and cell adhesion (Fig. 3B). These clustered terms were highly correlative with the GO terms of BP with high significance, showing the accuracy of identified changes in MSCs. Importantly, these results strongly suggested that angiogenesis and ECM organization (especially collagen fibril organization) were the primary cellular responses of MSCs to SF treatment, and individual GO terms associated with these two categories was shown in Fig. 3C. For CC and MF, the key GO terms associated with ECM, collagen, and growth factor were enriched, including “extracellular matrix”, “collagen type V trimer”, “platelet-derived growth factor binding”, and “growth factor activity” (Fig. S2A-B), indicating that the identified cellular changes in BP could be attributed to the secreted collagens and growth factors by MSCs, which was consistent with the observed changes in gene expression (Fig. 2J, 2L). Moreover, Kyoto Encyclopedia of Genes and Genomes (KEGG) enrichment analysis was conducted to recognize the activated signaling pathways of MSCs when exposed to SF. The results revealed several signaling pathways of MSCs to be potentially activated after SF stimulation, including PI3K-Akt, ECM-receptor interaction, focal adhesion, HIF-1, TGF-beta, and AMPK signaling pathways (Fig. 3D). SF or SF-based biomaterials has been reported to regulate cell behaviors by PI3K-Akt signaling pathway, and importantly this pathway is also found to mediate the collagen synthesis in mice and the angiogenic factor secretion in MSCs (*18, 54–56*). It well known that cell-matrix adhesion and interaction play an essentials role in various important biological processes, such as cell adhesion, cell proliferation, cell migration and regulation of gene expression (*57*). The up-regulated focal adhesion and ECM-receptor interaction indicated that the influence of SF on MSCs could be initiated by the interaction of SF and the membrane receptors of MSCs (Fig. 3D). Integrin family is the primary transmembrane adhesion receptors that connect ECM and cytoskeleton, playing a critical role in focal adhesion and ECM-receptor interaction. Thus, we examined the transcriptional levels of integrins in MSCs after SF treatment and found that SF significantly promoted the gene expression of various integrins, including *ITGA1*, *ITGA2*, *ITGA3*, *ITGA5*, *ITGB1*, and *ITGB3* (Fig. 3E). Interestingly, PI3K-AKT signaling is a downstream pathway of integrin signaling, and previous studies reported that the paracrine functions of MSCs could be enhanced by activating integrin/PI3K/AKT signaling pathways (*56*). Collectively, these findings highly suggested that SF triggered various cellular responses of MSCs by activating integrin signaling pathway. Among these changes, ECM organization and angiogenesis are the two most significant cellular events of MSCs in response to SF, which could be attributed to the increased expression of collagens and growth factors.

**Figure 3.**
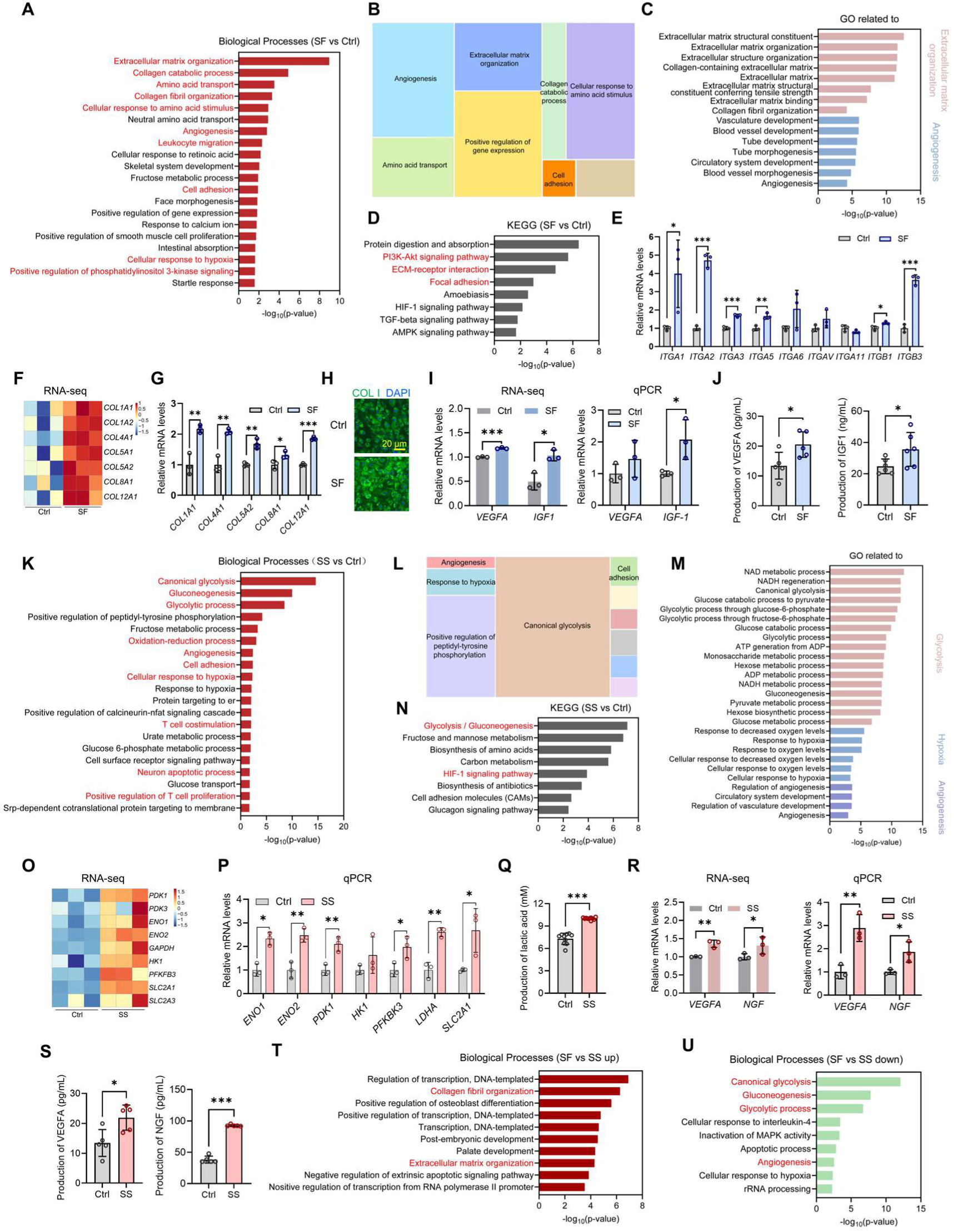
Functional insights of cellular changes of MSCs initiated by SF and SS. (A) Top GO terms of BP significantly enriched from up-regulated DEGs (SF vs Ctrl). (B) REVIGO analysis clusters all identified GO terms of BP into broader categories (SF vs Ctrl). (C) Significant GO terms associated with ECM organization and angiogenesis (SF vs Ctrl). (D) Top KEGG pathways significantly enriched from up-regulated DEGs (SF vs Ctrl). (E) The mRNA levels of integrin-related genes detected by qPCR. (F) Heatmap of expression levels of collagen-related genes by RNA-seq. (G) The mRNA levels of collagen-related genes detected by qPCR. (H) Representative images of IF staining for COL1 of MSCs with/without SF treatment. Scale bars = 20 μm. (I) The expression levels of *VEGFA* and *IGF-1* by RNA-seq and qPCR. (J) VEGFA and IGF-1 production of MSCs with/without SF treatment by ELISA. (K) Top GO terms of BP significantly enriched from up-regulated DEGs (SS vs Ctrl). (L) REVIGO analysis clusters all identified GO terms of BP into broader categories (SS vs Ctrl). (M) Significant GO terms associated with glycolysis, hypoxia, and angiogenesis (SS vs Ctrl). (N) Top KEGG pathways significantly enriched from up-regulated DEGs (SS vs Ctrl). (O) Heatmap of expression levels of glycolysis-related genes by RNA-seq. (P) The mRNA levels of glycolysis-related genes detected by qPCR. (Q) Lactic acid production of MSCs with/without SS treatment. (R) The expression levels of *VEGFA* and *NGF* by RNA-seq and qPCR. (S) VEGFA and NGF production of MSCs with/without SS treatment by ELISA. (T) Top GO terms of BP significantly enriched from up-regulated DEGs (SF vs SS). (U) Top GO terms of BP significantly enriched from down-regulated DEGs (SF vs SS). The results were presented as mean ± SD. *P<0.05, **P<0.01, ***P<0.001.

Therefore, we evaluated the expression of collagens and angiogenic growth factors in MSCs treated with SF. In transcriptional profiles of MSCs by RNA-seq, SF group exhibited a significantly higher expression of various collagens, such as *COL1A1*, *COL4A1*, *COL5A2*, *COL8A1*, and *COL12A1* as compared to the Ctrl group, which was subsequently validated by qPCR (Fig. 3F-G). Picrosirius red and immunofluorescence (IF) staining were performed to verify these changes of collagens at the protein level (Fig. 3H, S3). Picrosirius red staining and its quantitative analysis revealed that SF significantly promoted the collagen secretion and deposition of MSCs on days 3 and 7 as compared to the Ctrl group (Fig. S3). Besides, it was observed that SF group had a more intense staining for COL1 than the Ctrl group, indicating that an enhanced COL1 expression was induced by SF (Fig. 3H). These results demonstrated that SF significantly promoted the expression, secretion, and deposition of multiple collagens, which could contribute to the up-regulated ECM organization and collagen fibril organization of MSCs. In the past, the secreted ECM was generally considered as passive support for cells. However, recent studies have recognized it as a mediator of physical signals and an important regulator of various cell behaviors, including quiescence, proliferation, migration, and differentiation (*58, 59*). As the most abundant ECM constituents in various tissues bone, cartilage, skin, and gut, collagens have been found to play a critical role in regulating cell proliferation, migration, and differentiation (*60–63*). Thus, the notably increased collagen secretion in MSCs after SF treatment could influence ECM organization, thereby regulating the behaviors of themselves (autocrine) and surrounding cells (paracrine) in local environments for tissue repair and regeneration.

In addition to the changes in collagen secretion and ECM organization, MSCs treated with SF also exhibited a significantly enhanced capacity to promote angiogenesis. Angiogenesis is a complicated and multistep biological process that involves endothelial cell proliferation and migration, as well as the formation and stabilization of vessel structure, which is highly regulated by multiple factors, such as vascular endothelial growth factor (VEGF), insulin-like growth factor 1 (IGF-1), fibroblast growth factor 2 (FGF-2), nerve growth factor (NGF), and transforming growth factor beta (TGF-β) (*51, 64–66*). The results from RNA-seq revealed that the key angiogenic factors *VEGFA* and *IGF-1* were significantly up-regulated in the SF group compared with the Ctrl group, which was subsequently validated by qPCR (Fig. 3I). Enzyme-linked immunosorbent assay (ELISA) was performed to further evaluate the secretion of these angiogenic factors, and we found that MSCs in the SF group had a significantly higher production of VEGFA (1.61 times) and IGF-1 (1.44 times) than that of the Ctrl group (Fig. 3J). VEGFA is one of the most important angiogenic growth factors, and an increased VEGFA level can effectively induce the angiogenesis of epithelial cells and stem cells (*67–69*). Moreover, several studies have found that the CM from MSCs with a high level of VEGF and IGF-1 significantly increased blood vessel formation (*70, 71*). Previous studies also demonstrated that IGF-1 could promote angiogenesis via activating PI3K/Akt signaling pathway when stem cells and epithelial cells were co-cultured (*72*). Therefore, MSCs treated with SF could strongly support the angiogenesis of surrounding stem/progenitor cells and endothelial cells via paracrine secretion of VEGFA and IGF-1. In addition, it has been reported that IGF-1 promoted collagen expression and played a crucial role in regulating collagen remodeling (*73, 74*), possibly leading to increased collagen expression of MSCs in the SF group.

In summary, SF exhibited notable bioactivities to regulate multiple biological processes of MSCs, including ECM formation and organization, amino metabolism, angiogenesis, immunomodulation, and cell adhesion by activating integrin signaling. Among these cellular changes of MSCs, collagen-abundant ECM organization and angiogenesis were the most important cellular processes initiated by SF, which could largely be attributed to the increased secretion of a variety of collagens and angiogenic factors. Thus, SF was able to regulate the paracrine functions of MSCs and thereby formed a paracrine signaling network to guide the cellular behaviors of surrounding cells, thereby initiating multiple biological processes by regulating resident cells in local environment for tissue repair and regeneration.

#### 2.2.3 Functional insights of cellular changes of MSCs treated with SS

To investigate the cellular processes and pathways significantly affected by SS, GO enrichment analysis were also performed for up-regulated DEGs between SS and Ctrl. Focused on the GO terms with the smallest (strongest) P values, we found that multiple significant BP of MSCs were initiated by SS, including “canonical glycolysis”, “gluconeogenesis”, “glycolytic process”, “oxidation-reduction process”, “angiogenesis”, “cell adhesion”, “cellular response to hypoxia”, “T cell costimulation”, “neuron apoptotic process”, and “positive regulation of T cell proliferation” (Fig. 3K), which were associated with glycolysis, antioxidant activities, hypoxia adaptation, immunomodulation, and neuron survival. To further clarify the notable effects of SS on MSCs, REVIGO was employed to refine the numerous altered BP GO terms of MSCs. GO for BP were enriched to canonical glycolysis, positive regulation of peptidyl-tyrosine phosphorylation, response to hypoxia, angiogenesis, and cell adhesion (Fig. 3L). These results suggested that SS significantly promoted glycolysis, cellular response to hypoxia, and angiogenesis of MSCs, and individual GO terms associated with these three categories were presented in Fig. 3M. Besides, the key GO terms of CC and MF, such as “extracellular exosome”, “extracellular matrix”, and “growth factor activity” indicated that SS-induced cellular responses might also be mediated by the paracrine signals of MSCs (Fig. S4A-B). KEGG enrichment analysis confirmed that MSCs treated with SS had up-regulated glycolysis/gluconeogenesis and HIF-1 signaling pathway compared with the Ctrl group (Fig. 3N). Many studies have shown the close relationship of glycolysis and HIF-1 signaling that the activation of HIF-1 signaling pathway promotes the expression of various glycolytic enzymes and in turn the enhanced glycolysis also improves HIF-1α activity (*75*). Thus, we further evaluated the influence of SS on glycolysis and HIF-1 signaling pathway. It was observed that many glycolysis-related genes identified by RNA-seq, including *PDK1, PDK3, ENO1, ENO2, GAPDH, HK1, PFKFB3, SLC2A1*, and *SLC2A3*, were significantly up-regulated in the SS group as compared to the Ctrl group (Fig. 3O). We further validated the expression of these genes by qPCR and found consistent gene changes to those by RNA-seq (Fig. 3P). Moreover, we observed that MSCs in SS group exhibited a significantly increased *LDHA* expression and lactic acid production as compared to the Ctrl group, confirming that an enhanced glycolysis was initiated by SS stimulation (Fig. 3P-Q). Collectively, these results highly indicated that SS significantly promoted the metabolic switch of MSCs to glycolysis. Unfortunately, both the results of RNA-seq and qPCR showed no significant change in the expression of HIF-1 signaling pathway-related genes, including *HIF-1α*, *HIF-2α*, and HIF*-1β* on the designed time points (Fig. S5). Therefore, enhanced glycolysis rather than HIF-1 signaling is the main pathway by which SS regulates MSCs.

In addition, the enhanced angiogenesis seemed to be a critical cellular event of MSCs after SS treatments. We found that the angiogenic genes identified by RNA-seq, such as *VEGFA* and *NGF* were significantly up-regulated in the SS group as compared to the Ctrl group, which was subsequently validated by qPCR (Fig. 3R). Moreover, it was confirmed by ELISA that the secretion levels of VEGFA and NGF in the SS group were significantly higher than those in the Ctrl group (Fig. 3S). These finding demonstrated that SS significantly promoted angiogenic factor secretion of MSCs, which may contribute to the enhanced angiogenesis observed in the SS group (Fig. 3K-M). Interestingly, increasing evidence have shown an inseparable association between glycolysis and angiogenesis (*76–78*). Endothelial cells, the primary cell population for angiogenesis, were found to be highly glycolytic, and PFKFB3-driven glycolysis and followed glycolytic production of ATP could promote vessel sprouting by inducing the rearrangement of endothelial cells (*76, 77*). Besides, Liu et al. demonstrated that the highly glycolytic macrophages and microglia can enhance the expression of pro-angiogenic and pro-inflammatory cytokines, ultimately inducing pathological retinal angiogenesis (*78*). Notably, the highly glycolytic MSCs induced by SS also showed a significantly increased expression of angiogenic factors (e.g., VEGFA and NGF) and proinflammatory factors (e.g., IL-6, IL-32, and TNFAIP3) (Fig. 3O-R and S6). In addition, lactic acid, the primary product of glycolysis, has been shown to promote angiogenesis by up-regulating VEGF expression and secretion, so the increased lactic acid production of MSCs in the SS group may contribute to their enhanced angiogenesis (*79, 80*). Taken together, our results highly suggested that SS promoted the metabolic switch of MSCs to glycolysis, and the highly glycolytic MSCs could promoted angiogenesis by secreting more angiogenic factors.

In summary, SS regulated various biological processes of MSCs, including glucose metabolism, oxidative stress, angiogenesis, cell adhesion, adaptation to hypoxia, and immunomodulation. Among these cellular changes, glycolysis and angiogenesis were the most notable cellular processes in response to SS. Specifically, the highly glycolytic MSCs induced by SS are inclined to promote blood vessel formation by secreting various angiogenic factors rather than directing MSCs differentiation towards endothelial cells. Thus, these paracrine signals of MSCs stimulated by SS could be served as a strong angiogenic stimulus to induce angiogenesis by interacting with surrounding endothelial cells and stem/progenitor cells in the local environments.

After understanding the effects of SF and SS separately, we also compared the differences in the effects of SF and SS on MSCs. Key GO terms for BP with high significance showed that MSCs in the SF group had notably enhanced “collagen fibril organization”, and “extracellular matrix organization” while exhibited significantly down-regulated “canonical glycolysis”, “gluconeogenesis”, and “glycolytic process” compared with the SS group (Fig. 3T-U), which was consistent with the distinct effects of SF and SS (Fig. 3A-C and 3K-M). Although both SF and SS showed a capacity to promote angiogenesis by regulating the paracrine signals of MSCs, we observed that the SF group had down-regulated angiogenesis compared with SS group (Fig. 3U), indicating the stronger ability of SS to initiate angiogenic processes.

Taken together, although SF and SS are incapable of effectively guiding the directional differentiation of stem cells, they have an important influence on multiple biological processes and pathways of MSCs. Specifically, SF significantly enhanced ECM organization, collagen fibril organization, and angiogenesis of MSCs by secreting various collagens and angiogenic factors, with activation of integrin signaling pathway. SS effectively induced the metabolic switch of MSCs to glycolysis and subsequently promoted the secretion of angiogenic factors to induce angiogenesis. Interestingly, notable changes were found in the genes encoding secreted proteins of MSCs upon SF or SS treatment, strongly suggesting that SF and SS initiated these cellular events by dominantly regulating paracrine signals of MSCs.

### 2.3 Cellular secretome changes of MSCs triggered by SF and SS

We next focused on the paracrine products of MSCs in response to SF and SS. Proteomic analysis was performed to investigate the secretome changes of MSCs treated with/without SF or SS treatment for 3 days, as illustrated in Fig. 4A. Principal component analysis (PCA) of proteomic data for cellular secretome revealed that the replicates of each group assembled into a clear cluster and showed similar features (Fig. 3B). As shown by the Venn plot, MSCs shared a highly similar secretome among the three groups, with a total of 206 identical proteins recognized, indicating that both SF and SS mainly induced the changes of cellular secretome in quantity rather than type (Fig. 4C). Specifically, we identified 47 differentially expressed proteins (DEPs) (P<0.05) in SF vs Ctrl, with 35 up-regulated proteins and 12 down-regulated proteins (Fig. 4D). In contrast, we recognized that 10 proteins were significantly up-regulated while 48 proteins were significantly down-regulated in SS vs Ctrl (Fig. 4E). When SF group was compared to SS group, 46 DEPs were up-regulated and 11 DEPs were down-regulated (Fig. 4F). GO and KEGG enrichment analysis were conducted to obtain the functional insights into these secretome changes (Fig. 4G-J). Notably, several BP GO terms associated with skin development and wound healing were enriched. In SF vs Ctrl, we identified several key GO terms specific to “epidermis development”, “keratinization”, “cell adhesion”, “skin development”, “keratinocyte differentiation”, “positive regulation of epidermis development”, “peripheral nervous system axon regeneration”, “epithelial cell differentiation”, and “wound healing” (Fig. 4G). Similarly, several GO terms for BP were significantly enriched in SS vs Ctrl, including “epidermis development”, “extracellular matrix organization”, “cell adhesion’, “collagen fibril organization”, “keratinization”, “glycolytic process”, “regulation of cell adhesion”, “regulation of cell growth”, “epithelial cell differentiation”, and “wound healing” (Fig. 4H). The above results strongly suggested that the paracrine signals provided by MSCs in response to SF and SS have potentials to regulate skin repair and regeneration. Subsequently, KEGG analysis revealed that the changed secretome in the SF group were associated with multiple cellular pathways, including “ECM-receptor interaction”, “PI3K-Akt signaling pathway”, “focal adhesion”, and “complement and coagulation cascade” (Fig. 4I). In addition to these pathways, “fructose and mannose metabolism”, “HIF-1 signaling pathway”, and “glycolysis/gluconeogenesis” were enriched from significantly changed paracrine products of MSCs in SS vs Ctrl (Fig. 4J). These signaling pathways enriched from changed secretome of MSCs after SF or SS treatment exhibited a high degree of consistency with our RNA-seq data (Fig. 3D, 3N), confirming the critical roles of these paracrine signals in the cellular responses to SF and SS. Since the RNA-seq results showed significant changes in genes associated with growth factors and collagen-related ECM, we specially investigated the secretion of their corresponding proteins (Fig. 4K). Consistent with our RNA-seq data, significant changes in the expression and secretion of these proteins were observed, and the SF groups particularly had a significantly higher secretion of collagens (COL1A1, COL1A2, COL4A1, COL5A2, COL6A1, COL6A3, and COL11A1) and growth factors (IGFBP3, IGFBP4, and IGFBP7) compared to the Ctrl or SS group (Fig. 4K).

**Figure 4.**
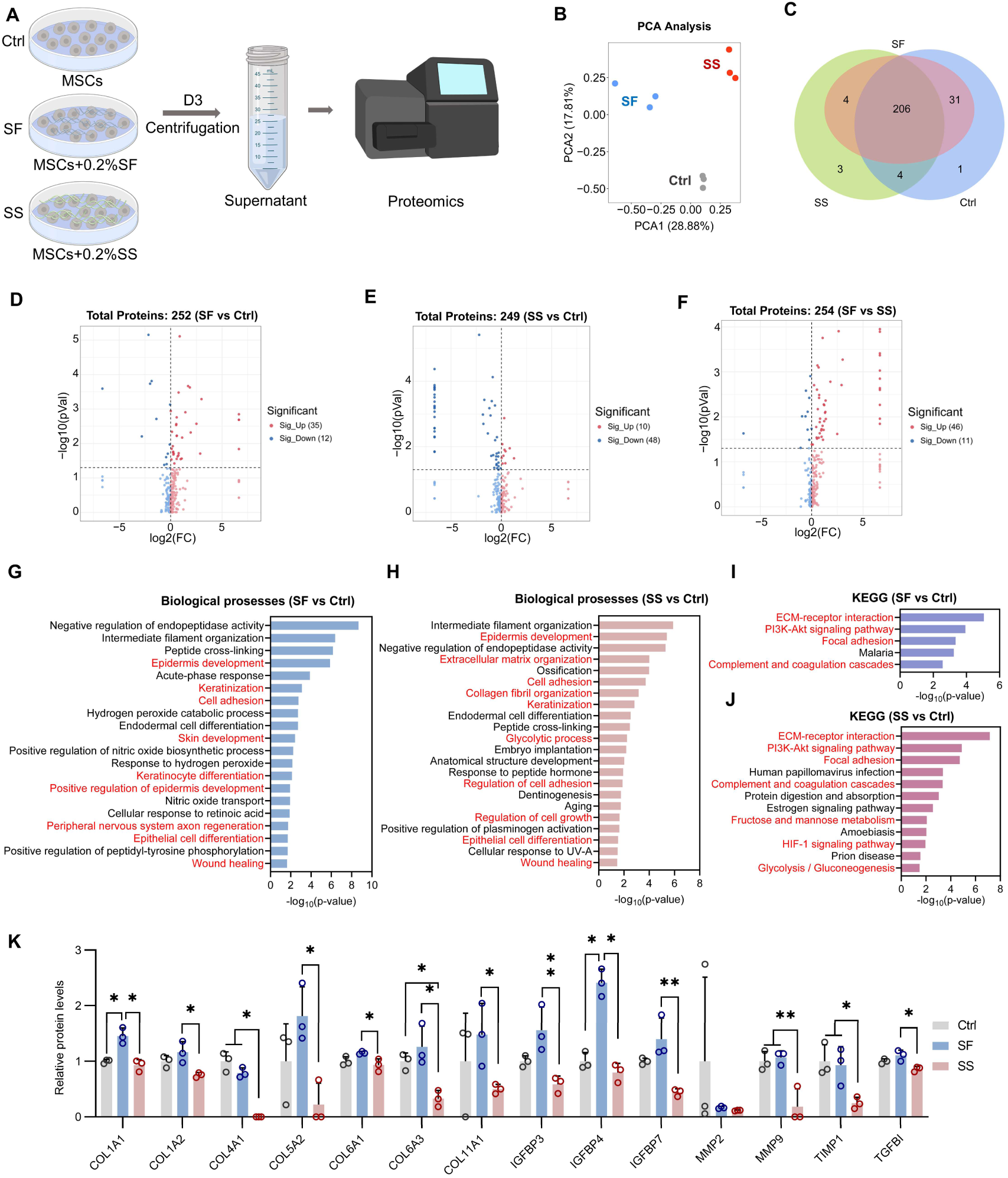
Proteomics reveals cellular secretome changes initiated by SF and SS. (A) Schematic of the proteomic profiling of the secreted proteins from MSCs treated with/without SF or SS. (B) PCA plot of proteomic data in different groups. (C) Venn plot of identified proteins in the supernatant. (D-F) Volcano plots of secreted protein levels in SF vs Ctrl, SS vs Ctrl, and SF vs SS groups. (G-H) Top GO terms of BP significantly enriched from all DEPs in SF vs Ctrl and SS vs Ctrl groups. (I-J) Top KEGG pathways significantly enriched from all DEPs in SF vs Ctrl and SS vs Ctrl groups. (K) The protein levels of secreted factors associated with collagens, growth factors, and matrix proteases. The results were presented as mean ± SD. *P<0.05, **P<0.01.

In general, both SF and SS significantly influenced the paracrine productions of MSCs, which played a critical role in the biological processes and signaling pathways they triggered. Moreover, consistent with our RNA-seq data and in vitro verification, the secretome of MSCs initiated by SF or SS showed great potentials to regulate wound healing, especially the repair and regeneration of skin.

### 2.4 Paracrine signals of MSCs triggered by SF and SS regulate resident cellular behaviors of skin wound healing in vitro

Our integrated transcriptomics and proteomics had revealed that both SF and SS play significant roles in promoting wound healing-related biological processes, such as angiogenesis, collagen synthesis, and immunomodulation by regulating the paracrine signals of MSCs, thereby showing great potentials for skin repair and regeneration. We therefore reasoned that the paracrine signals of the transplanted MSCs in response to SF or SS could strongly regulate multiple processes of wound healing by influencing the behaviors of the resident cells in the microenvironments of impaired skin. Skin wound healing is mediated by multiple biological processes, including inflammation, angiogenesis, contractile granulation formation, and ECM deposition, which involve various cell types, such as immune cells, endothelial cells, and fibroblasts (*81*). Thus, we investigated whether the paracrine factors of MSCs treated with SF or SS significantly regulated the cellular behaviors of these resident cells involved in skin wound healing. To probe the effects of paracrine products of MSCs, the CMs of the MSCs treated with/without SF or SS were collected and applied to culture fibroblasts, endothelial cells, and macrophages, which is illustrated in Fig. 5A.

**Figure 5.**
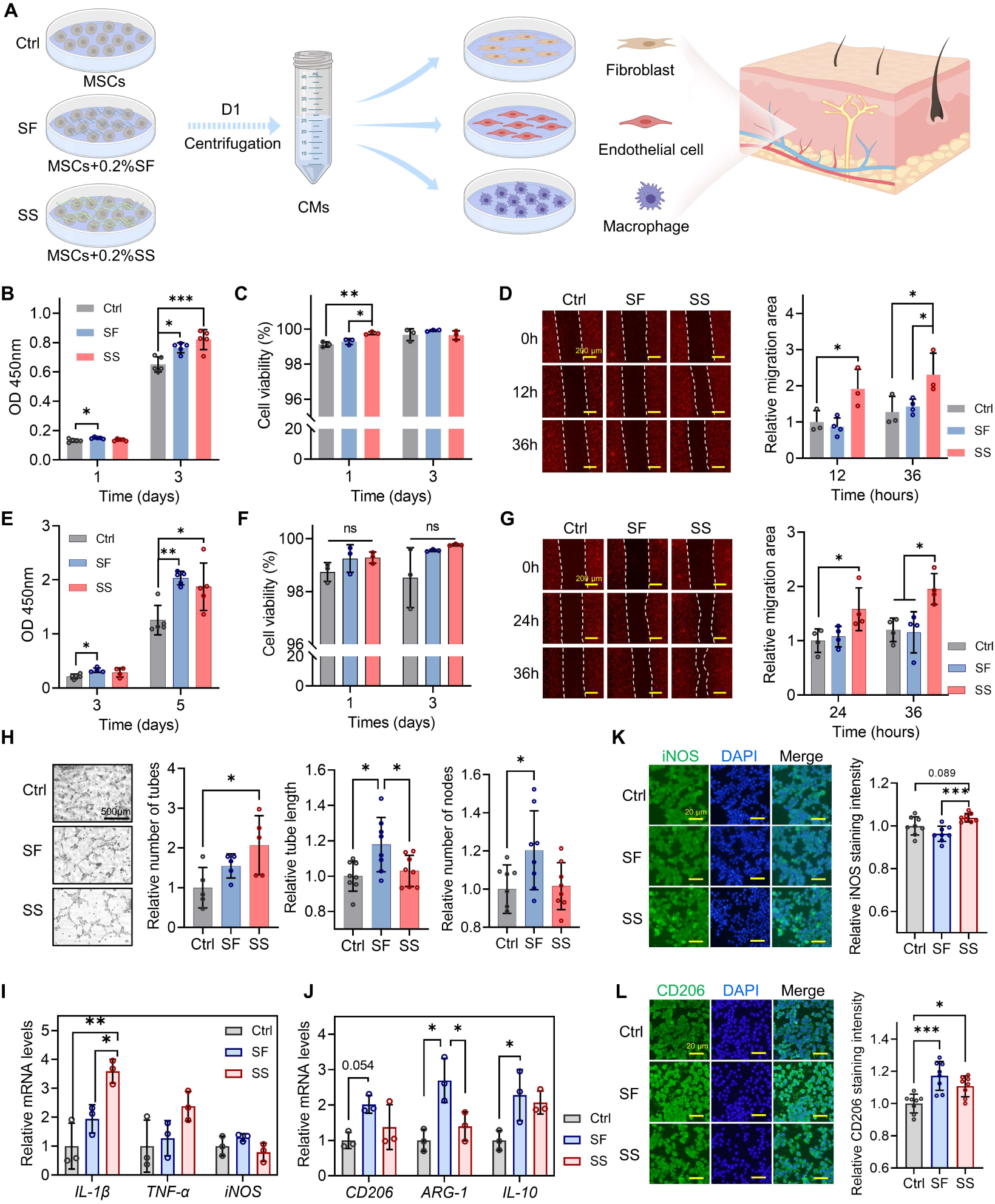
Paracrine signals of MSCs stimulated by SF and SS regulate resident cellular behaviors of wound healing. (A) Schematic of the in vitro cell culture experiments for evaluating the effects of paracrine signals of MSCs on wound healing. (B) Fibroblasts proliferation with different CM treatments measured by CCK-8. CMs were collected from MSCs treated with/without SF or SS. (C) Cell viability of fibroblasts treated with different CMs by live/dead staining. (D) Cell migration of fibroblasts with different CM treatments using scratch assay. Scale bars = 200 μm. (E) HUVECs proliferation with different CM treatments measured by CCK-8. (F) Cell viability of HUVECs treated with different CMs by live/dead staining. (G) Cell migration of HUVECs with different CM treatments using scratch assay. Scale bars = 200 μm. (H) Representative images and quantitative analysis of tube formation of HUVECs with different CM treatments for 6 h. Scale bars = 500 μm. (I-J) The gene expression levels of pro-inflammatory markers (*IL-1β, TNF-α*, and *iNOS*) and anti-inflammatory markers (*CD206, ARG-1*, and *IL-10*) of macrophages treated with different CMs for 1 day. (K-L) Representative images and quantitative analysis of IF staining for iNOS and CD206 treated with different CMs for 1 day. Scale bars = 20 μm.The results were presented as mean ± SD. *P<0.05, **P<0.01, ***P<0.001.

#### 2.4.1 Paracrine signals of MSCs in response to SF and SS regulate the proliferation, viability and migration of fibroblasts

Fibroblasts are mesenchymal cells with multiple functions including ECM deposition, regulation of epithelial differentiation, and immunomodulation, playing a dominant role in wound healing (*82*). At the end of inflammatory phase, fibroblasts migrate to the wound area, and subsequently proliferate and mediate the fibrin clot degradation, ECM deposition, as well as wound contraction, ultimately initiating the proliferative phase of wound healing (*82*).Thus, to evaluate the potentials of the paracrine signals from MSCs for skin repair, its effects on fibroblast behaviors were firstly investigated. CCK-8 assay revealed that fibroblasts treated with both SF CM and SS CM exhibited a significantly increased proliferation rate on days 1 and/or 3 as compared to the Ctrl CM and there was no significant difference between SF and SS groups, indicating the positive role of these paracrine signals of MSCs induced by SF or SS in promoting fibroblasts proliferation (Fig. 5B). Besides, it was observed that the CM collected from the MSCs treated with SF and SS slightly increased the cell viability of fibroblasts as compared to the Ctrl CM on day 1 (Fig. 5C). Subsequently, we evaluated the effects of different CMs on fibroblasts migration via a scratch assay. Representative images and quantitative analysis revealed that the SS CM significantly promoted the migration of fibroblasts compared to that of the Ctrl CM and SF CM, while the SF CM had no obvious pro-migratory effect on fibroblasts (Fig. 5D). Taken together, these findings suggested that the secretion of paracrine factors from the MSCs induced by SF could promote fibroblasts proliferation while the signals stimulated by SS enhanced both the proliferation and migration of fibroblasts.

To explore the mechanisms underlying the relationship between the paracrine signals from MSCs and fibroblasts behaviors, we found that IGF-1 has been reported to enhance the viability and proliferation of fibroblasts (*83, 84*). Thus, the increased IGF-1 secretion of MSCs in response to SF could contribute to the improved viability and proliferation of fibroblasts in the SF group (Fig. 3J). Besides, NGF has also been found to promote the proliferation and migration of fibroblasts (*85*), and a significantly increased secretion of NGF was detected in the CM obtained from the SS group (Fig. 3S). Additionally, it has been widely reported that FGF can promote the proliferation and migration of fibroblasts (*86*). Our results of RNA-seq revealed that MSCs in both SF and SS groups exhibited a higher expression level of *FGF2* than that of the Ctrl group (Fig. S7). Moreover, it was observed that MSCs in the SS group had a significantly increased *FGF11* expression compared to the Ctrl and SF groups (Fig. S7). Thus, the increased FGF expression of MSCs in response to SF or SS could also contribute to the enhanced fibroblast proliferation and/or migration. Taken together, these increased growth factors in the CMs collected from the MSCs treated with SF or SS could enhance fibroblasts proliferation and migration and might be beneficial for wound healing and skin regeneration in vivo (*87, 88*).

#### 2.4.2 Paracrine signals of MSCs in response to SF and SS regulate the proliferation, migration, and tube formation of endothelial cells

Angiogenesis is recognized as a crucial activity during wound healing as it provides vascular supply to support cells with oxygen and nutrition in damaged tissue (*89*). It is a complex biological process that involves multiple cellular events, including the proliferation and migration of endothelial cells, as well as the formation and stabilization of vessel structure, which are regulated by a variety of growth factors, such as VEGF, IGF-1, FGF-2, and NGF (*51, 64–66*). Due to the increased expression and secretion of these angiogenic factors, we reasoned that MSCs treated with SF or SS could promote the angiogenic processes in a paracrine manner. Thus, proliferation, migration, and tube formation assays were performed to confirm the effects of these paracrine signals on HUVECs. CCK-8 assay revealed that HUVECs showed a significantly enhanced proliferation when cultured with the CM obtained from the MSCs treated with SF or SS as compared to the Ctrl CM, and no significant difference was identified between the SF CM and SS CM (Fig. 5E). However, the CM collected from the MSCs exposed to SF or SS did not significantly influence the cell viability of HUVECs (Fig. 5F). Subsequently, the effects of these paracrine signals on cell migration were evaluated through a scratch assay. After a culture of 24 or 36 h, only the CM from MSCs treated with SS could enhance the migration of HUVECs, while the SF CM showed minimal effects (Fig. 5G). Finally, when HUVECs were exposed to different CMs for 6 h, SF CM showed a significantly improvement of tube formation with regards to tube length and node number as compared to the Ctrl and/or SS group, while SS CM increased the number of tubes compared with the Ctrl group (Fig. 5H). VEGF is a dominant angiogenic mediator and it can initiate angiogenic cascades by activating its various components (*90*). Our results showed that both SF and SS significantly promoted the secretion of VEGFA from MSCs (Fig. 3J, 3S), thereby leading to the enhanced angiogenesis of HUVECs treated with SF and SS CMs. Moreover, the increased levels of IGF-1 and NGF in CMs from MSCs treated with SF and SS respectively have also been reported to induce angiogenesis of endothelial cells (*91, 92*). Taken together, these results indicated that the secretion of paracrine factors of MSCs in response both to SF and SS promoted the proliferation and tube formation of HUVECs, and the SS-induced paracrine signals was also able to enhance the migration HUVECs, suggesting the great potentials of these signals to promote angiogenesis during wound healing.

#### 2.4.3 Paracrine signals of MSCs in response to SF and SS regulate the polarization of macrophages

Inflammation is an early response mediated by various immune cells to eliminate cellular debris, foreign body, and bacteria during wound healing (*93*). Macrophage is one of the earliest immune cell types to arrive at the skin wounds and mediate an acute pro-inflammatory response that often lasts for a few days (*93*). After that, the pro-inflammatory (M1) phenotype of macrophages will switch towards an anti-inflammatory (M2) phenotype, which can promote inflammation resolution, tissue remodeling, and new vessel formation to accelerate wound healing (*93*). However, the malfunction of macrophage phenotypic switch will result in chronic inflammation in some diseases (e.g., diabetes), which inhibited the healing processes of wounds (*89*). Due to the effects of SF and SS on the immunomodulation of MSCs, we investigated the influence of their paracrine products on macrophage polarization by qPCR and IF staining. qPCR was performed to investigate the expression of pro-inflammatory (*IL-1β*, *TNF-α*, and *iNOS*) and anti-inflammatory (*CD206*, *ARG-1*, and *IL-10*) markers of macrophages when cultured with different CMs for 1 day. Our results revealed that SS CM significantly up-regulated the expression of *IL-1β* as compared to that of the Ctrl and SF CMs (Fig. 5I). The gene expression of *TNF-α* exhibited a similar trend to that of *IL-1β* while no significance was detected between the three groups (Fig. 5I). Besides, macrophages in the SF group had a substantially increased expression of *CD206*, *ARG-1*, and *IL-10* as compared to the other two groups (Fig. 5J). IF staining further showed that SS CM promoted the protein expression of iNOS and CD206 while SF CM only significantly increased the CD206 expression of macrophages (Fig. 5K-L). Generally, these results indicated that the paracrine signals of MSCs stimulated by SF promoted the macrophage polarization towards anti-inflammatory phenotype while that stimulated by SS was more inclined to activate inflammation of macrophages with up-regulation of both pro- and anti-inflammatory markers.

To better understand the macrophage polarization induced by paracrine signals of MSCs, we investigated the expression of inflammation-related genes in MSCs after SF and SS treatments. The results revealed that various inflammation-related genes were differentially expressed between Ctrl, SF, and SS groups, including *CXCL12*, *IL-6*, *IL-16*, *IL-32*, *TNFAIP3*, *TNFSF9*, and *TNFSF15* (Fig. S6). Notably, the SS group exhibited a significantly increased expression of pro-inflammatory markers *(IL-6*, *IL-32*, and *TNFAIP3)* and a significantly reduced expression of anti-inflammatory markers *(CXCL12* and *TNFSF9)* in MSCs as compared to the Ctrl and SF groups, and these combined changes have been reported to induce pro-inflammatory responses and macrophage polarization towards M1 phenotype (*94–100*). In contrast, the majority of these pro- or anti-inflammatory factors were not significantly changed between the SF and Ctrl groups, while the SF group exhibited a reduced TNFSF15 expression compared to the Ctrl group, which could facilitate macrophage polarization towards M2 phenotype (Fig. S6) (*101*). It was worth mentioning that these changed inflammatory factors exhibited similar, different, or even inverse effects on macrophage phenotypes, so the macrophage polarization mediated by paracrine signals of MSCs was more inclined to be determined by the combined effects of these factors. Taken together, our results suggested that the paracrine signals of MSCs stimulated by SF and SS can induce macrophage polarization into anti-inflammatory and pro-inflammatory phenotypes, respectively, which could play a role in regulating inflammation resolution or activation during wound healing. From the perspective of immunomodulation, SF is more suitable than SS for skin wound healing due to the anti-inflammatory responses of MSCs to SF, especially for diseases with chronic inflammation, such as diabetes.

In general, our findings showed that the paracrine products of MSCs, stimulated by SF or SS, could effectively regulate the behaviors of resident cells in skin microenvironments, including promoting proliferation, viability, and migration of fibroblasts, enhancing angiogenesis of endothelial cells, as well as modulating M1/M2 polarization of macrophages, showing great promises of SF/SS-mediated MSC-based therapies for wound healing.

### 2.5 SF and SS modulate the paracrine function of MSCs to promote skin wound healing in vivo

Due to the notable effects of SF- or SS-triggered paracrine signals of MSCs on promoting various healing processes in vitro, we speculated that the use of SF or SS as the carrier material for MSCs had the potential to improve skin wound healing in vivo. To ensure that these paracrine signals could be produced and secreted in vivo, here we used Gelatin methacryloyl (GelMA) hydrogel as the base material of SF and SS for the encapsulation of MSCs, providing a suitable three-dimensional (3D) environment for the biochemical interactions of SF and SS with MSCs. GelMA hydrogel was able to support MSCs survival owing to its good biocompatibility and water-rich environment, and could be photocrosslinked in situ and adhere well to injured skin (*102*). By using this hydrogel system, we created a highly physiological 3D environment for the survival and growth of MSCs, and ensured the sustained release of paracrine signals of MSCs in response to SF or SS, ultimately applying to the treatment of full-thickness skin defects in rat.

#### 2.5.1 Preparation of hydrogel system for MSC-SF/SS interactions in 3D environment

As illustrated in Fig. 6A-B, MSC-seeded photocrosslinked GelMA hydrogels incorporated with/without 0.2% (w/v) SF or SS were successfully fabricated, termed as Ctrl (GelMA hydrogel), SF (SF-GelMA hydrogel) and SS (SS-GelMA hydrogel). The incorporation of SF or SS did not significantly alter the physical properties of GelMA hydrogels, including surface morphology, pore size, porosity, and rheological behaviors (Fig. 6C-F). Therefore, it ensured that the changes in paracrine function and cellular behavior of MSCs were solely affected by the biochemical signals from SF or SS, but not by their physical properties. Then, live/dead staining was performed to evaluate the cell viability of MSCs encapsulated in the 3D hydrogels, and results demonstrated that MSCs in each group had a high cell viability of >92%, showing no significant difference between three groups (Fig. 6G). Subsequently, to verify the capacity of SF and SS to regulate the paracrine functions of MSCs in the 3D environment, mass spectrometry imaging (MSI), picrosirius red staining and ELISA were performed to detect cell metabolites and secretions of MSCs interacting with SF or SS in the hydrogel system. MSI revealed that both SF and SS in 3D hydrogels notably increased the production of cell metabolites compared to the Ctrl group, and MSCs treated with 0.2% (w/v) SF for 3 days exhibited the highest levels of cell metabolites compared to the other concentrations or other culture periods (Fig. 6H, S8). Besides, SF significantly enhanced collagen production of MSCs as compared to Ctrl and SS, as evidenced by picrosirius red staining on sections of 3D hydrogels (Fig. 6I). For angiogenic factors, SF and SS exhibited significantly increased IGF-1 and VEGFA secretion, respectively, in 3D hydrogels compared to the Ctrl group (Fig. 6J-K), showing a similar trend to two-dimensional (2D) cultures (Fig. 2J, 3J, 3S). Consistent with 2D cultures, it was also observed in 3D hydrogels that the SS-GelMA hydrogel had a higher NGF production than the Ctrl group, indicating the potential of SS to modulate neurogenesis (Fig. 6L, 3S). Besides, SS significantly enhanced the production of lactic acid when MSCs were cultured in SS-GelMA hydrogels, suggesting enhanced glycolysis of MSCs after SS treatments in both 2D and 3D cultures (Fig. 6M, 3O-Q). Taken together, these data confirmed that SF and SS could promote the paracrine function of MSCs in our developed 3D hydrogel system with high concordance with 2D cultures, enhancing their translational potential in the treatment of skin defects in vivo.

**Figure 6.**
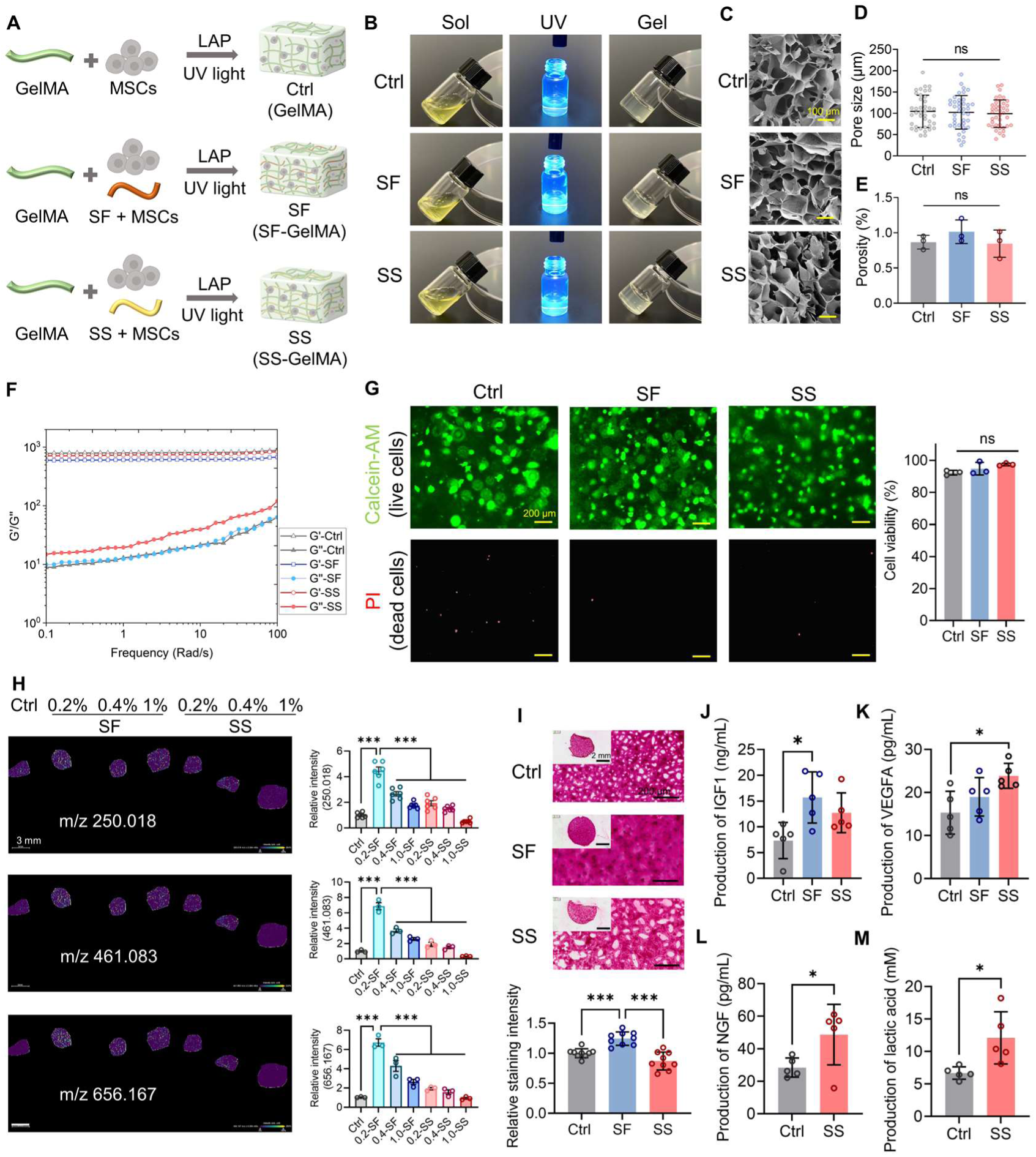
Preparation and evaluation of 3D hydrogel system for MSC-SF/SS interactions. (A) Schematic of the preparation of MSC-seeded photocrosslinked GelMA hydrogels incorporated with/without 0.2% (w/v) SF or SS. (B) Sol-gel transition of GelMA, SF-GelMA and SS-GelMA hydrogels. (C) SEM images of three hydrogels. Scale bars = 100 μm. (D-E) Pore size and porosity of three hydrogels. (F) Rheological frequency sweeps of three hydrogels. (G) Representative images and quantitative analysis of live/dead staining of MSCs in 3D hydrogels of different groups for 3 days. Scale bars = 200 μm. (H) Spatial distribution and quantitative analysis of cell metabolites of MSCs in 3D hydrogels for 3 days by MSI. Scale bars = 3 mm. (I) Representative images and quantitative analysis of picrosirius red staining of MSCs-seeded 3D hydrogels. Low magnification: scale bars = 2 mm; high magnification: scale bars = 200 μm. (J-L) Production of IGF-1, VEGFA and NGF of MSCs in 3D hydrogels by ELISA. (M) Production of lactic acid of MSCs in Ctrl and SS hydrogels. The results were presented as mean ± SD. *P<0.05.

#### 2.5.2 Paracrine signals of MSCs triggered by SF and SS promote skin repair and regeneration

As illustrated in Fig. 7A, rat full-thickness skin defect model was established, and the developed 3D hydrogels were applied to evaluate the effects of paracrine signals of MSCs interacting with SF or SS on wound healing in vivo. Gross observation of skin wounds with different treatments on days 0, 3, 7, 10, and 14 were shown in Fig. 7B. Compared with the Ctrl group, both SF and SS groups exhibited a significantly accelerated wound healing on days 7 and 10 (Fig. 7B-C). Specifically, both the SF (59.83%) and SS (62.90%) groups achieved about 60% wound healing while the Ctrl group showed only 33.54% wound closure on day 7 (Fig. 7C). Similarly, we found that on day 10, the wound healing rate was close to 90% in the SF (89.68%) and SS (90.37%) groups, while that in the Ctrl group was only 74.11% (Fig. 6C). No significant difference was identified in the wound healing rate between the SF and SS groups, indicating the comparable effect of SF and SS on MSCs in accelerating wound healing. Histological evaluation using hematoxylin and eosin (H&E) and Masson’s Trichrome (MT) staining was performed to assess the formation and maturation of repaired tissue on day 14 (Fig. 7D-E). It was observed that all three groups had complete epidermal regeneration and well-formed and integrated collagen-like tissue formation in the dermis area. However, evident skin appendages, characterizing the complete remodeling of dermis, were presented in the SF and SS groups, but not in the Ctrl group (Fig. 7D-E). Quantitative analysis for skin thickness revealed that both the SF (∼1.86 mm) and SS (∼1.88 mm) groups has a significantly increased skin thickness compared to the Ctrl group (∼1.46 mm), while no significant difference was identified between the Ctrl and SS group (Fig. 7F). Besides, SF and SS enhanced angiogenesis of newly-formed skin tissue, as evidenced by the significantly higher number and larger cross-sectional area of the neovessels than those in the Ctrl group (Fig. 7G-H), which might be attributed to the higher secretion of angiogenic factors from MSCs in SF and SS groups (Fig. 6J-M). Additionally, a higher number of hair follicles was observed in the SF (∼17.27/mm^2^) and SS (∼11.03/mm^2^) groups as compared to the Ctrl group (∼4.80/mm^2^) (Fig. 7I). These results demonstrated that the SF and SS groups induced an enhanced skin repair and regeneration structurally and functionally.

**Figure 7.**
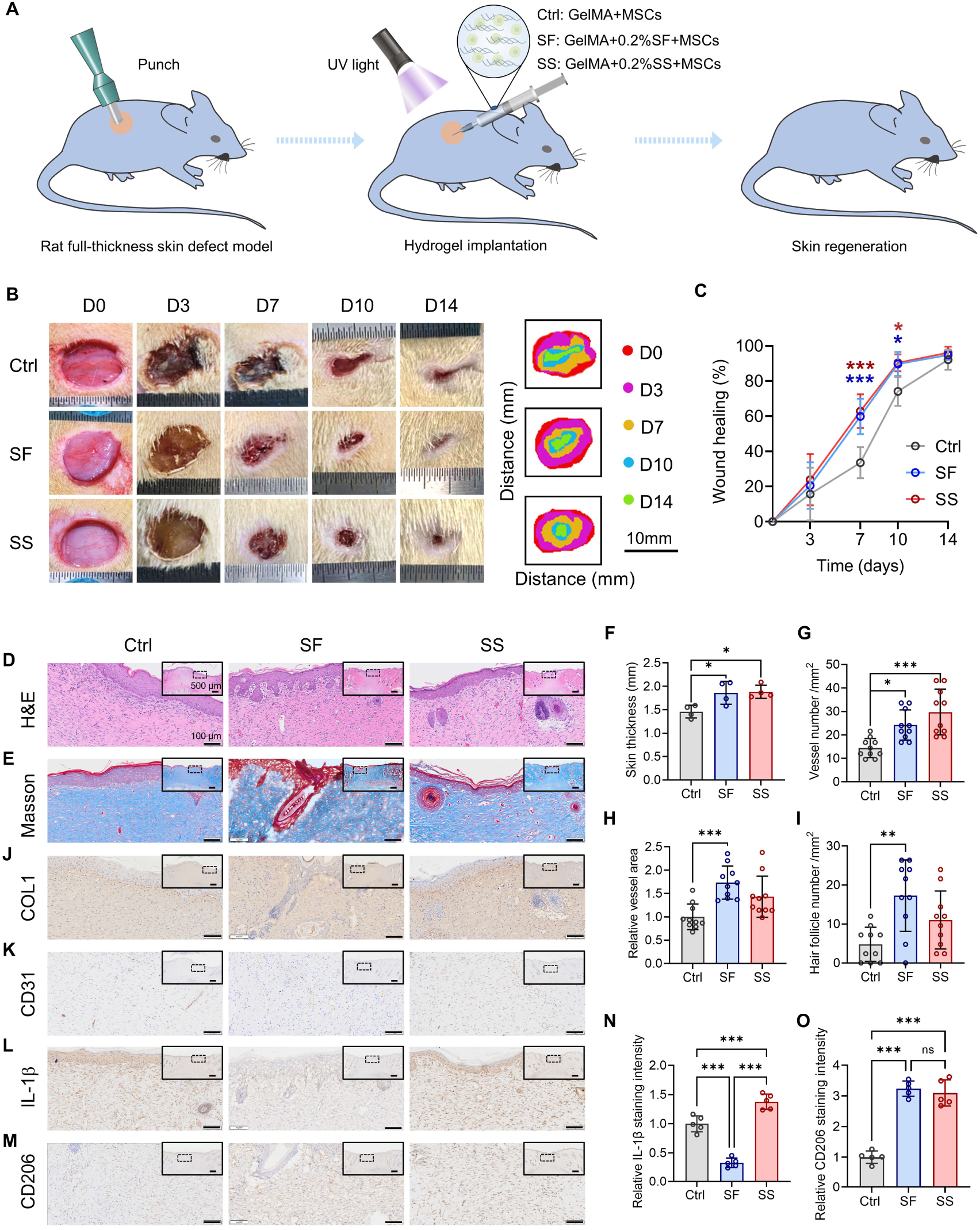
Paracrine signals of MSCs triggered by SF and SS promote skin repair and regeneration in vivo. (A) Schematic of the animal study procedure. MSC-seeded GelMA hydrogels incorporated with/without 0.2% (w/v) SF or SS were applied to each wound. (B) Gross morphology of the wound healing process within 14 days. (C) Quantitative analysis of wound healing rate in different groups. (D-E) H&E and MT staining of regenerated skin tissues in different groups on day 14. Low magnification: scale bars = 500 μm; High magnification: scale bars = 100 μm. (F-I) Quantitative analysis of skin thickness, vessel number, vessel area, and hair follicle number from H&E stained images. (J-K) Representative images of IHC staining for COL1 and CD31 in different groups. Low magnification: scale bars = 500 μm; High magnification: scale bars = 100 μm. (L-O) Representative images and quantitative analysis of IHC staining for IL-1β and CD206 in different groups. Low magnification: scale bars = 500μm; High magnification: scale bars = 100 μm. The results were presented as mean ± SD. *P<0.05, **P<0.01, ***P<0.001.

It is worth mentioning that both SF and SS biomaterials by themselves have been reported to promote wound healing (*16, 103, 104*). To reveal the contribution of the paracrine signals of MSCs triggered by SF and SS to wound healing, and to exclude the intrinsic effects of SF and SS, unseeded hydrogels were also applied to treat rat skin defects (Fig. S9A). Different from the results obtained with MSC-seeded hydrogels, the unseeded SF-GelMA or SS-GelMA hydrogel did not significantly enhance wound healing rate at each time point, as compared to the unseeded GelMA hydrogel (Fig. S9B-C). Moreover, H&E staining showed that only the vessel number was increased in the SS group as compared to the Ctrl and SF groups (Fig. S9D-E). We further compared the healing rate and tissue maturation of the wounds upon SF and SS treatments with/without MSCs. The results revealed that both SF and SS treatments with MSCs significantly improved wound closure as compared to that without MSCs on days 7 and 10 (Fig. S10-S11). In the SF or SS group with MSCs, the number of newly-formed blood vessels and hair follicles was substantially higher than that without MSCs (Fig. S12), which may be related to the increased levels of angiogenic factors secreted by MSCs upon SF/SS stimulation, such as VEGFA, IGF-1, and NGF (Fig. 6J-L). Collectively, these results confirmed the critical role of MSCs in wound healing and strongly suggested that the paracrine signals of MSCs stimulated by SF and SS, rather than SF and SS themselves, contributed to the improved skin regeneration in our study. Subsequently, histological and immunohistochemical (IHC) staining for COL1, CD31, IL-1β, and CD206 was performed to evaluate collagen composition, vessel formation, and inflammation level of the regenerated tissues in different groups. As shown in Fig. 7J, an intense staining for COL1 was observed in all groups, and both the SF and SS groups had a slightly stronger staining than that in the Ctrl group (Fig. 7J). COL1 was the most abundant collagen in the normal skin, and the higher COL1 deposition in the SF and SS group indicated the formation of more mature skin tissues (*105*). The staining for CD31, a maker of vascular endothelial cells, confirmed that an enhanced angiogenesis was initiated in the SF and SS groups (Fig. 7K), which was consistent with the quantified analysis from H&E images (Fig. 7G-H). Besides, the staining for IL-1β and CD206, as well as the quantitative analyses revealed that the SF group exhibited a significantly decreased staining intensity of IL-1β while a notably increased staining intensity of CD206 (Fig. 7L-O), suggesting that the paracrine signals of MSCs triggered by SF could inhibit inflammatory responses and promote macrophage polarization towards M2 phenotype, thereby facilitating functional regeneration of skin (*89*). In contrast, the SS group had a more intense staining of both IL-1β and CD206 than the Ctrl group (Fig. 7L-O), which was consistent with the in vitro results (Fig. 5I-L). Due to the simultaneous increase of pro- and anti-inflammatory markers, therefore, it was insufficient to clarify the polarization status of macrophages but supported a higher inflammatory level in the SS group.

Taken together, our results demonstrated that the paracrine signals of MSCs stimulated by SF and SS significantly promoted the repair and regeneration of skin with both structural and functional recovery. Compared with SF, SS induced a stronger inflammatory response in the wounds because of their distinct effects on immunomodulation.

#### 2.5.3 Global assessments of the wound microenvironments after treatment by proteomics

To globally evaluate the wound microenvironments after different treatments, proteomic analysis was conducted on the tissue samples (n=3/group) collected from the regenerated skin in rats on day 14 (Fig. 8A). PCA plot showed that the samples of each group assembled into a cluster, indicating a high consistency of the replicates from one group (Fig. S13). The heatmap of all identified proteins after cluster analysis exhibited a notable difference in the protein components between Ctrl, SF, and SS groups (Fig. 8B). Then, the pairwise comparisons of Ctrl, SF, and SS groups were performed to identify the DEPs (P<0.05). Specifically, we found that 78 proteins were differentially expressed between SF and Ctrl groups, with 47 up-regulated proteins and 31 down-regulated proteins (Fig. 8C). As for the SS and Ctrl groups, we detected 79 DEPs totally, and 38 of them were up-regulated and 41 of them were down-regulated (Fig. 8D). Besides, 15 up-regulated DEPs and 32 down-regulated DEPs were identified when the SF group was compared to the SS group (Fig. 8E). Among these DEPs, 39 and 58 DEPs were specific to SF vs Ctrl and SS vs Ctrl, and only 9 DEPs were specific to SF vs SS (Fig. 8F). Fig. 8G-H exhibited the hub 20 DEPs that were screened by CytoHubba in Cytoscape, which could reveal the more important changes in the wound environments after different treatments. Notably, we found that the SF group had significantly higher expression of ECM-related proteins including COL1A1, COL2A1, DCN, and TIMM13 compared to the Ctrl group, while the SS group showed a notably increased expression of glycolysis-related proteins, including GPI, PGAM2, PGK1, ALDOA, ENO2, ENO3, and GAPDH in wound environments than the Ctrl group, which was consistent with the results from in vitro studies (Fig. 3E-F, 3O-P). These results indicated that the increased collagen-enriched ECM deposition of MSCs induced by SF and the enhanced metabolic switch to glycolysis of MSCs triggered by SS could promote the healing process of wounds.

**Figure 8.**
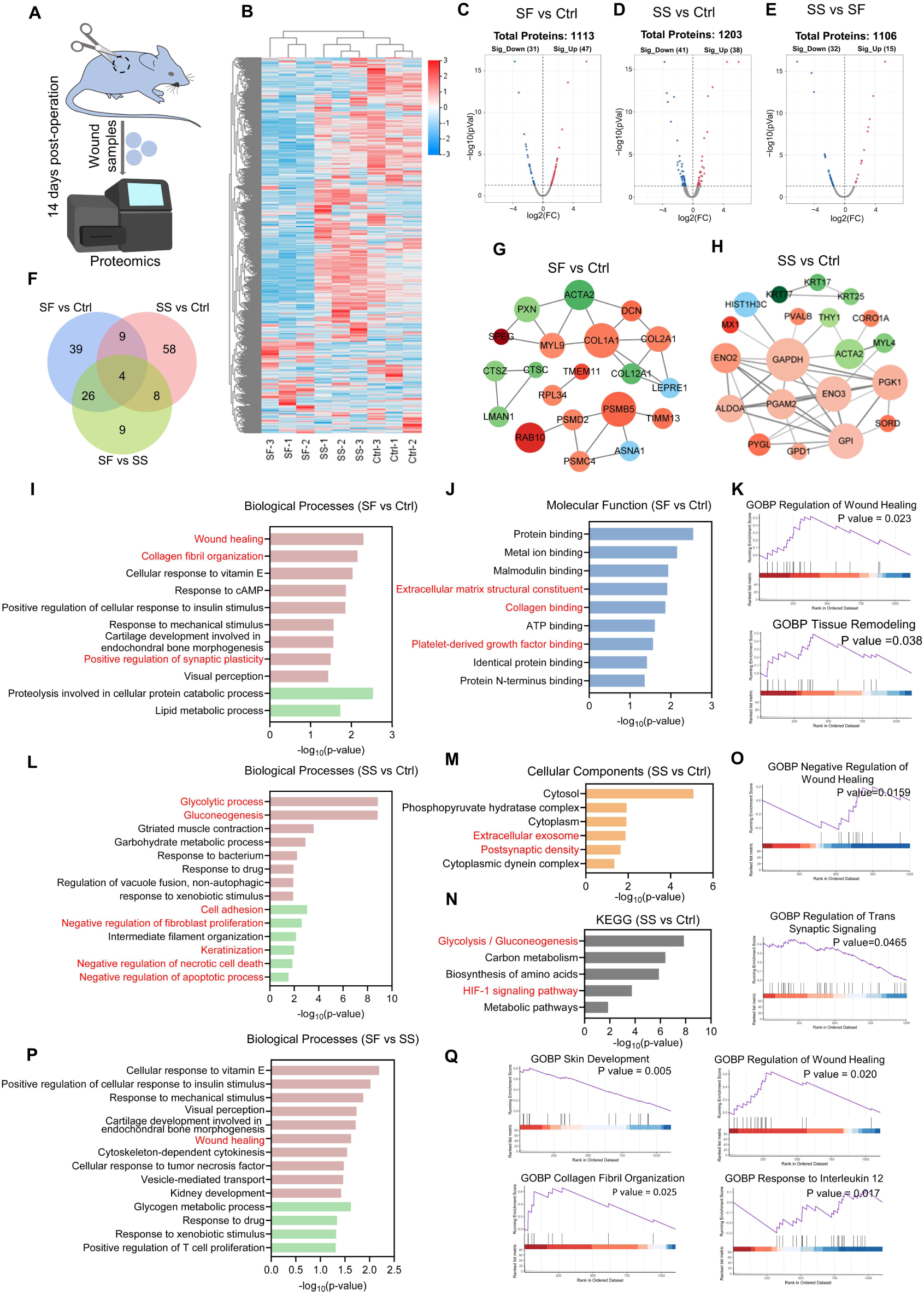
Global assessments of the wound microenvironments after treatment by proteomics. (A) Schematic of the proteomic profiling of the expressed proteins in wound environments. (B) Heatmap of protein expression levels after hierarchical cluster analysis. (C-E) Volcano plots of protein expression levels in SF vs Ctrl, SS vs Ctrl, and SF vs SS groups. (F) Venn plot of identified DEPs in pairwise comparison. (G-H) Protein interaction networks of top 20 hub DEPs in SF vs Ctrl and SS vs Ctrl groups. A larger size of protein node represents a higher frequency in the protein networks. The up-regulated proteins are presented in red circle while the down-regulated proteins are in green circle. (I) Significant GO terms for BP enriched from up-regulated (red) and down-regulated (green) DEPs (SF vs Ctrl). (J) Significant GO terms for MF enriched from up-regulated DEPs (SF vs Ctrl). (K) GSEA plots of cellular processes associated with wound healing (SF vs Ctrl). (L) Significant GO terms for BP enriched from up-regulated (red) and down-regulated (green) DEPs (SS vs Ctrl). (M) Significant GO terms for CC enriched from up-regulated DEPs (SS vs Ctrl). (N) Significant KEGG pathways enriched from up-regulated DEPs (SS vs Ctrl). (O) GSEA plots of cellular processes associated with wound healing (SS vs Ctrl). (P) Significant GO terms for BP enriched from up-regulated (red) and down-regulated (green) DEPs (SF vs SS). (Q) GSEA plots of cellular processes associated with wound healing (SF vs SS).

To further obtain the functional insights in these DEPs of the wound microenvironments, GO enrichment analyses were subsequently conducted. In SF vs SS, several BP GO terms were significantly enriched from up-regulated DEPs, including “wound healing”, “collagen fibril organization”, and “positive regulation of synaptic plasticity” (Fig. 8I). Besides, the SF group presented several significant up-regulated GO terms for CC and MF, such as “extracellular matrix”, “extracellular matrix structural constituent”, “collagen binding”, and “platelet-derived growth factor binding” as compared to the Ctrl group (Fig. 8J, S14A), which could probably be mediated by the paracrine signals of MSCs. Moreover, gene set enrichment analysis (GSEA) was performed to overcome the limitations of DEPs analysis, revealing that SF-treated MSCs activated the “regulation of wound healing”, “tissue remodeling”, and “axon guidance”, while suppressed “leukocyte chemotaxis” during wound healing, leading to the enhanced skin regeneration in the SF group as compared to the Ctrl group (Fig. 8K, S14B). Specifically, the up-regulated “positive regulation of synaptic plasticity” and “axon guidance” by SF treatment might be attributed to the increased NGF expression of MSCs in response to SF (Fig. S15).

In terms of SS vs Ctrl, we identified a notably enhanced “glycolytic process”, “gluconeogenesis”, and “posphopyruvate hydratase activity” among the significant GO terms in BP and MF (Fig. 8L, S16A), indicating the enhanced glycolysis in the wounds of SS groups, which was also confirmed by KEGG and GSEA analyses (Fig. 8N, S16B). Besides, we also observed some down-regulated BPs that contributed to the accelerated wound healing, including “negative regulation of fibroblast proliferation”, “negative regulation of necrotic cell death”, and “negative regulation of apoptotic processes” (Fig. 8L). Significant GO terms for CC showed that “extracellular exosome” and “postsynaptic density” were up-regulated (Fig. 8M), indicating the improved transmission of paracrine and neural signals during wound healing. In addition, it was observed that HIF-1 signaling was activated in the SS group as compared to the Ctrl group, which probably initiated the metabolic switch to glycolysis (Fig. 8N). Consistent with the increased NGF secretion in vitro and the accelerated wound healing in vivo, GSEA analysis revealed that the SS group exhibited a positive regulation of transsynaptic signaling and wound healing (Fig. 6L, 7C, 8O).

Finally, we also compared the differences between SF and SS groups. Although similar wound healing effects was shown by gross observation and histological evaluation, we discovered from the in vivo proteomics that the SF group was more beneficial to wound healing due to the enhanced “skin development”, “wound healing”, “regulation of wound healing”, “collagen fibril organization”, and “epithelial cell differentiation” as well as the decreased “response to interleukin 12” in SF vs SS comparisons, as evidence by the GO and GSEA analysis (Fig. 8P, 8Q, S17).

Taken together, our in vivo results demonstrated that SF and SS effectively promoted skin repair and regeneration by potentiating the paracrine signals of MSCs. The paracrine signals of MSCs triggered by SF or SS could influence the behavior of multiple resident cells in skin wound microenvironments and regulate key processes of skin wound healing, including inflammation, angiogenesis, fibroblast activities, and ECM deposition. Although both SF and SS are capable of regulating the paracrine functions of MSCs and improving wound healing, their underlying mechanisms are distinct (Fig. 9). SF mainly promoted the secretion of IGF-1, VEGFA, collagens, and more anti-inflammatory factors from MSCs to induce cell proliferation, enhance angiogenesis and modulate macrophage polarization towards M2 phenotype via activating integrin signaling. SS could induce the secretion of VEGFA and NGF and ultimately enhance cell proliferation, migration, angiogenesis and inflammatory responses of macrophages by promoting the metabolic switch of MSCs to glycolysis. The higher pro-inflammatory responses induced by SS may impair the repair and regeneration of skin defects with chronic inflammation. Therefore, in this regard, SF is more suitable than SS as a carrier material for MSCs for skin wound healing.

**Figure 9.**
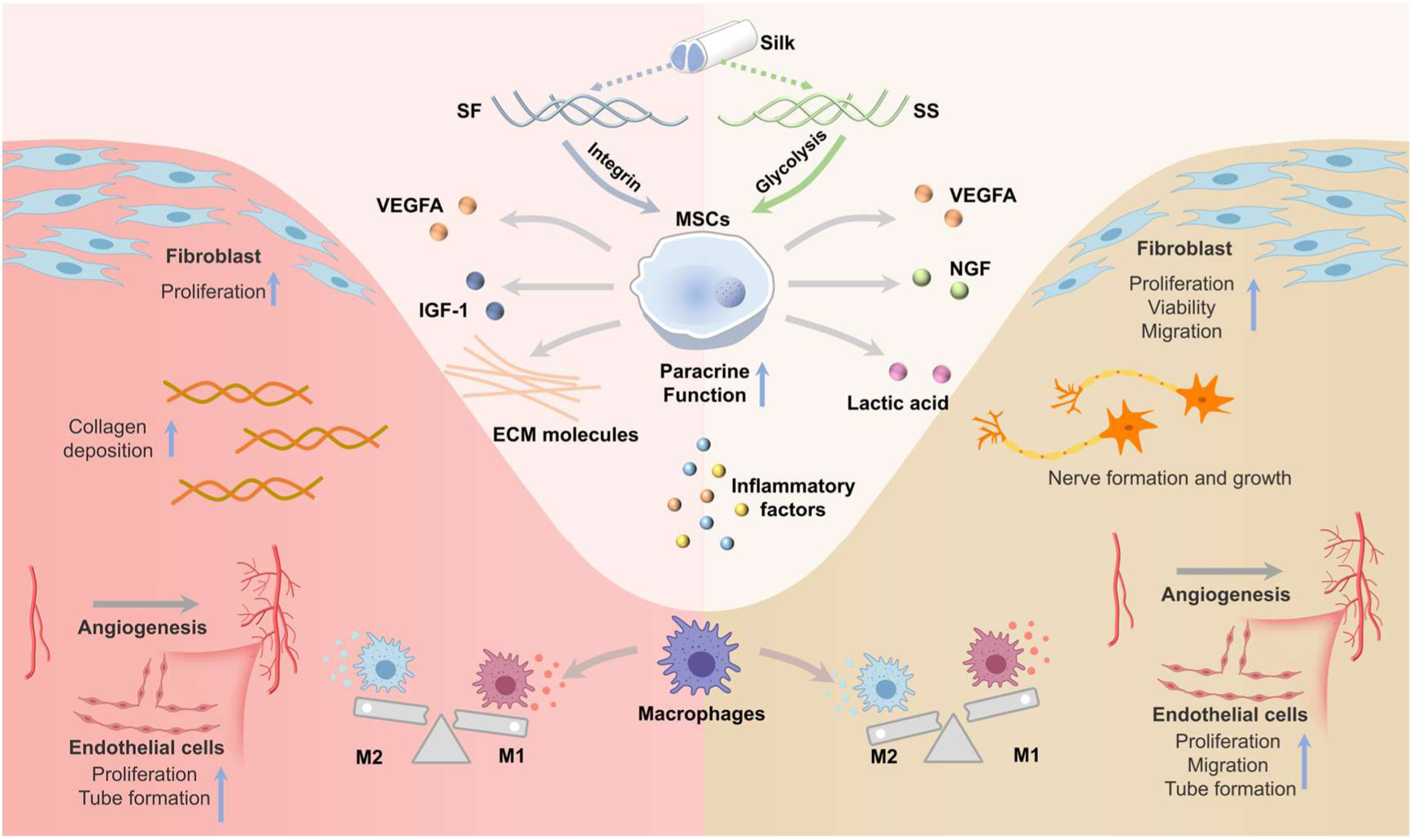
Schematic illustration of SF and SS promoting skin wound healing by potentiating the paracrine signals of MSCs.

## 3. Conclusions

In this study, we applied multiomics to comprehensively understand the biochemical interactions of SF and SS with MSCs, as well as the followed changes in biological processes and molecular activities. Integrated RNA-seq and proteomic analysis for transcriptional profiles and cellular secretome consistently demonstrated that SF and SS initiated widespread and strong cellular responses, and notably potentiated the paracrine activities of MSCs regulating ECM deposition, angiogenesis, and immunomodulation via differentially activating integrin and glycolysis signaling pathways, rather than directly determining stem cell fate. These paracrine signals of MSCs stimulated by SF and SS effectively improved skin repair and regeneration by influencing the behavior of multiple resident cells in skin wound microenvironments. From the perspective of immunomodulation, SF seemed to be a more suitable than SS as a carrier material for MSCs for skin wound healing due to its better anti-inflammatory effects. To our knowledge, this is the first study to provide global and reliable insights into the cellular interactions with SF and SS via multiomics analysis, which contributes directly to the design of silk-based scaffolds for tissue engineering and stem cell therapy. Furthermore, our findings highlight the importance and feasibility of “omics” techniques for the biological evaluation of cell-biomaterials interactions, which may have board implications for future development of tissue engineering scaffolds in terms of material selection and scaffold design.

## 4. Materials and methods

### Materials and methods

#### Preparation and characterization of SF and SS

SF and SS were prepared as described previously (*106, 107*). Briefly, for SF preparation, raw silk fibers (*Bombyx mori*, Zhejiang Xingyue Biotechnology, Hangzhou, China) were boiled in 0.02 M Na2CO3 (Sinopharm, China) for 30 min to remove the outer SS of the fibers. The degummed fibers were washed with Milli-Q water, dried at 37 ℃, and then dissolved in 9.3 M LiBr (Aladdin, Shanghai, China) for 4 h at 60 °C. The dissolved SF solution were added into a dialysis bag (3500 MWCO, Yuanye, China) and dialyzed against the Milli-Q water for 48 h (six changes of water) to remove LiBr. The dialyzed SF solution was centrifugated at 9000 rpm for 20 min at 4 °C twice. Finally, the supernatant was collected and the concentration of the SF solution was determined. For SS preparation, raw silk fibers were washed with Milli-Q water and then boiled in 0.02 M Na2CO3 (Sinopharm, China) for 1 h. After removing the silk fibers, the remained SS solution was collected and added into a dialysis bag (3500 MWCO, Yuanye, Shanghai, China) and dialyzed against the Milli-Q water for 48 h (six changes of water). The SS solution was subsequently dialyzed against 20% (w/v) polyethylene glycol (PEG, 10,000 MW, Sinopharm, Beijing, China) solution for 48 h to obtain the concentrated SS solution. Finally, the SS solution was collected and its concentration was determined.

#### Cell culture

All types of cells were cultured in the humidified atmosphere with a 5% CO2 at 37°C. Primary human MSCs (Cyagen Biosciences, Suzhou, China) from bone marrow were cultured in a minimum essential medium alpha (MEM-α, Gibco, Carlsbad, CA) medium with 10% fetal bovine serum (FBS, Wisent, Canada), and 1% Penicillin-Streptomycin (P/S, Gibco, Carlsbad, CA). The MSCs between passages 3 to 8 were used in the following experiments. L929 fibroblast (a gift from Prof. Jie Chao of Southeast University, China), human umbilical vein endothelial cells (HUVEC, a gift from Prof. Jie Chao of Southeast University, China), and RAW 264.7 macrophages (a gift from Prof. Lixin Wang of Southeast University, China) were cultured in a high-glucose (4.5g·L^-1^) Dulbecco’s Modified Eagles’ Medium (DMEM, Gibco, Carlsbad, CA) with a supplementary of 10% FBS and 1% P/S. The medium was changed every three days.

For the collection of conditioned medium (CM), MSCs were treated with/without 0.2 % (w/v) SF and SS in the FBS-free medium. After a culture of 1 day, the supernatant of the medium was collected and then centrifuged at 2000 rpm to remove dead cells and cell debris. The supernatant of centrifuged medium was collected and stored at -20 °C. When used for cell culture, CM was mixed with the complete growth medium at a ratio of 1:1.

#### Cell proliferation

Cell proliferation was evaluated by cell counting kit-8 (CCK-8, APExBIO, Houston, TX). Cells were seeded into 96-well plates at the density of 2×10^3^ cells/well and cultured in fresh growth medium. MSCs were treated with SF and SS in various concentrations for 1, 3, and 7 days to determine an appropriate concentration for cell proliferation. Fibroblasts or HUVECs were treated with/without 0.2% (w/v) SF or SS for 1 and 3 days. At the designated time points, the medium was replaced with a 10% (v/v) CCK-8 solution. After an incubation of 1 h at 37 °C, the absorbance of the incubated solution was determined by using an 800 TS microplate reader (BioTek, Winooski, VT, USA) at 450 nm.

#### Cell viability and metabolic activity

Cell viability was assessed by live/dead staining. In two-dimensional (2D) culture, 2×10^3^ cells were seeded into 96-well plates and cultured in fresh growth medium. MSCs, fibroblasts, and HUVECs were treated with/without 0.2% (w/v) SF or SS for 1-7 days. In three-dimensional (3D) culture, 2×10^5^ MSCs were encapsulated in Gelatin methacryloyl (GelMA) hydrogels and cultured in fresh growth medium for 3 days. At the designated time points, the samples in 2D or 3D culture were incubated in 50ul or 500 µl Calcein-AM/PI Double Staining Kit (Dojindo, Japan) working solution at 37 °C for 20 min, respectively. Subsequently, the live and dead cells of the samples were observed and imaged using fluorescence microscopy (Carl-Zeiss, Oberkochen, Germany). The cell viability was determined by calculating the ratio of live cells to total cells.

The Alamar Blue assay was used to evaluate the metabolic activity of MSCs. 2 ×10^3^ MSCs were seeded into 96-well plates and cultured in fresh growth medium. Then, MSCs were treated with/without 0.2% (w/v) SF or SS for 1 and 3 days. The culture medium was replaced with a growth medium with 10% (v/v) Alamar blue solution at the designated time points. After incubated for 2 h, the fluorescence intensity of the dye was measured at 590 nm after an excitation wavelength of 560 nm on a Varioskan™ LUX multimode microplate reader (Thermo Fisher Scientific, Madison, WI).

#### F-actin staining

Cell morphology and adhesion were visualized by F-actin staining. 1×10^3^ MSCs were seeded into 96-well plates and cultured in fresh growth medium with/without 0.2% (w/v) SF or SS for 7 days. On day 7, the samples were fixed in 4% paraformaldehyde (PFA) for 10 min, and then immersed in 0.1% (v/v) TritonX-100 (Beyotime, Beijing, China) for 5 min to enhance cell permeability. Subsequently, the samples were incubated in the Actin-Tracker Green (Beyotime, Beijing, China) work solution with a dilution of 1:100 for 1 h. The DAPI (Beyotime, Beijing, China) with a dilution of 1:2000 was used for nuclear staining for 5 min. The stained samples were observed and imaged by a fluorescence microscope (Carl-Zeiss, Oberkochen, Germany). The relative staining intensity of the sample was measured using ImageJ software (NIH, Bethesda, MD, USA).

#### Cell migration

Cell migration was evaluated by a scratch test. 5×10^4^ MSCs, fibroblasts, or HUVECs that were pre-stained with Dil (Beyotime, Beijing, China) were seeded into 24-well plates for 90% confluence and cultured in fresh growth medium. On the next day, a 1-ml pipet tip was used to make a scratch in the center of the confluent cell monolayers, and cells were gently rinsed with PBS twice. Subsequently, cells were treated with growth medium with/without 0.2% (w/v) SF or SS, or the different CMs for 12-36 h. The samples were collected at 0, 12, 24, and 36 h and imaged by a fluorescence microscope (Carl-Zeiss, Oberkochen, Germany). The changes in the scratched area were quantified using ImageJ software (NIH, Bethesda, MD) and the relative migration area was calculated.

#### Tube formation assay

A tube formation assay was performed to assess the angiogenesis of HUVECs. 10 ul Matrigel (Corning, Bebford, MA) was added to the pre-cooled ibidi μ-Slide angiogenesis dish (ibidi, Munich, Germany) and allowed to solidify at 37°C for 30 min. Then, 5×10^3^ HUVECs were seeded in the prepared wells and cultured in fresh growth medium with/without 0.2% (w/v) SF or SS treatment. After culture for 6 h, the tube formation of HUVECs was observed and imaged using an optical microscope (Olympus, Tokyo, Japan). The quantitative analyses for tube number, tube length, and node number were performed respectively.

#### Osteogenic, adipogenic, and chondrogenic differentiation

Osteogenic differentiation: 1×10^4^ MSCs were seeded into 24-well plates and cultured with the osteoinductive medium consisting of high-glucose DMEM medium, 10% FBS, 1% P/S, 10 mM b-glycerol phosphate (Sigma, St. Louis, MO), 10 nM dexamethasone (Sigma, St. Louis, MO), and 50 μg/ml ascorbic acid (Sigma, St. Louis, MO). MSCs were cultured with an osteoinductive medium with/without 0.2% (w/v) SF or SS for 7 days, and the medium was changed every two days. The osteogenic differentiation of MSCs was examined by using BCIP/NBT Alkaline Phosphatase Color Development Kit (Beyotime, Beijing, China). The samples were observed and imaged using a light microscope (Olympus, Tokyo, Japan). The staining intensity (IntDen/Area) was measured using ImageJ software (NIH, Bethesda, MD).

Adipogenic differentiation: 1×10^4^ MSCs were seeded into 24-well plates and were cultured in high glucose DMEM medium with 10% FBS, 1% P/S, 1 mM dexamethasone, 10 μg/ml insulin (Sigma, St. Louis, MO), and 0.5 mM isobutylxanthine (Sigma, St. Louis, MO). MSCs were treated with/without 0.2% (w/v) SF or SS in the adipogenic differentiation medium for 14 days, and the medium was changed every two days. Adipogenic differentiation of MSCs was evaluated by lipid droplets staining with 0.6% (w/v) oil red O (Solarbio, Beijing, China). The stained samples were observed and imaged using a light microscope (Olympus, Japan). For quantitative analysis, the staining was destained with 100% isopropyl alcohol (Sinopharm, Beijing, China), and the absorbance of the extracted dye was measured at 510 nm on a Varioskan™ LUX multimode microplate reader (Thermo Fisher Scientific, Madison, WI).

Chondrogenic differentiation: 5×10^3^ MSCs were seeded into 96-well plates and cultured with chondrogenic medium consisting of high-glucose DMEM medium, 1% Insulin-TransferrinSelenium (ITS, Gibco, Carlsbad, CA), 1 mM sodium pyruvate (Gibco, Carlsbad, CA), 50 μg/ ml ascorbic acid (Sigma, St. Louis, MO), and 10 ng/ml TGF-β1 (PeproTech, Rocky Hill, NJ) in the presence of 0.2% (w/v) SF or SS for 14 days. At the designated time point, alcian blue staining (1%, pH = 2.5, Macklin, Shanghai, China) was performed to examine glycosaminoglycan (GAG) deposition for chondrogenesis evaluation. After gross examination using a light microscope (Olympus, Tokyo, Japan), the staining of samples was destained with 6 M Guanidine-HCl (Aladdin, Shanghai, China) overnight. The absorbance of the extracted dye was determined by using an 800 TS microplate reader (BioTek, Winooski, VT) at 630 nm.

#### Picrosirius red staining

The synthesis and deposition of collagens were evaluated by picrosirius red staining. In 2D culture, 1×10^4^ MSCs were seeded into 24-well plates, and cultured in growth medium with/without 0.2% (w/v) SF for 3 and 7 days. In 3D culture, 2×10^5^ MSCs were encapsulated in GelMA hydrogels with/without 0.2% (w/v) SF or SS and cultured in the fresh growth medium for 7 days. At the designated time points, the samples in 2D or 3D culture were stained with picrosirius red (G-Clone, Beijing, China) for 1 h. The stained samples were observed and imaged by using a stereo-microscope (SZ61, Olympus, Japan). For quantitative analysis, the staining was destained with 0.1M NaOH (Sinopharm, Beijing, China), and the absorbance of the extracted dye was measured at 540 nm on a Varioskan™ LUX multimode microplate reader (Thermo Fisher Scientific, Madison, WI).

#### RNA isolation and qPCR

Total RNA was isolated from cultured cells according to the protocols of manufacturers (Tiangen, Beijing, China). cDNA was subsequently synthesized by using a cDNA reverse transcription kit (Toyobo, Japan), and gene expression was examined by real-time PCR using primer pairs (Genscript, Nanjing, China) and SYBR Green (Invitrogen, Waltham, MA). The primers used in this study were presented in Table S1. GAPDH or β-actin was selected as a housekeeping gene.

#### RNA-seq and bioinformatics analysis

MSCs were cultured in growth medium with/without 0.2% (w/v) SF or SS (n = 3/group). After a culture of 3 days, cells were collected and analyzed by using a high-output HiSeq platform. Gene expression levels were normalized by calculating the fragments per kilobase of transcript per million reads (FPKM). The transcriptomic data was analyzed on the DAVID website (https://david.ncifcrf.gov/), OmicStudio tools (https://www.omicstudio.cn/), and REVIGO website (http://revigo.irb.hr/). After paired comparison of identified genes, the differentially expressed genes (DEGs) with a P value less than 0.05 were recognized and used for subsequent Gene Ontology (GO) and Kyoto Encyclopedia of Genes and Genomes (KEGG) enrichment analyses on the DAVID website. REVIGO was employed to reduce redundant GO terms. RNA-seq datasets were available in the Sequence Read Archive (SRA) with the accession number SRP388527.

#### Proteomic analysis for the secretome of MSCs

MSCs were cultured in growth medium with/without 0.2% (w/v) SF or SS (n = 3/group). After a culture of 3 days, the supernatant of cultured MSCs in each group was collected and centrifuged at 2000 rpm to remove dead cells and cell debris. Proteomic analysis based on the inbuilt label-free quantification (LFQ) was conducted to examine to secretome of MSCs after SF or SS treatment. The transcriptomic data was analyzed on the DAVID website (https://david.ncifcrf.gov/) and OmicStudio tools (https://www.omicstudio.cn/). The paired comparison between three groups was performed to recognize the secretome changes of MSCs. The differentially expressed proteins (DEPs) with a P value less than 0.05 and were recognized and used for GO and KEGG enrichment analyses. The mass spectrometry proteomics data have been deposited to the ProteomeXchange Consortium (http://proteomecentral.proteomexchange.org) via the iProX partner repository with the dataset identifier PXD035603.

#### Enzyme-linked immunosorbent assay (ELISA)

MSCs were seeded into 24-well plates (2×10^4^ /well) or 200 μl GelMA hydrogels (4×10^4^ /hydrogel) and immersed in the growth medium. After being cultured with/without 0.2% (w/v) SF or SS for 3 days, the supernatant of MSCs was collected. ELISA were performed to detect the levels of VEGFA, IGF-1, and NGF in the supernatant according to the manufacturers’ instructions (Proteinetch, Wuhan, China). The production of lactic acid in the supernatant was detected using a Lactic Acid Assay Kit (Jiancheng, Nanjing, China).

#### Immunofluorescence (IF) staining

MSCs and macrophages were seeded into 96-well plates at a density of 1×10^3^. MSCs were cultured in growth medium with/without 0.2% (w/v) SF or SS for 7 days, and macrophages were cultured with different CMs from MSCs for 1 day. IF staining for COL1 in MSCs, as well as iNOS and CD206 in macrophages, was performed. Briefly, cells were fixed in 4% PFA for 20 min, permeabilized with 1% (v/v) Triton X-100 (Beyotime, Beijing, China) for 10 min, and blocked with QuickBlock™ Blocking Buffer for Immunol Staining (Beyotime, Beijing, China) for 20 min, successively. Subsequently, samples were incubated with rabbit anti-COL1 (Proteintech, Wuhan, China), anti-iNOS (Affinity Biosciences, OH, USA), and anti-CD206 (Proteintech, Wuhan, China) at 4 °C overnight. After being rinsed with PBS, the samples were stained by 488-conjugated goat anti-rabbit IgG (Proteintech, Wuhan, China) for 1 h at room temperature. DAPI was used for nuclear staining for 5 min. The staining of samples was observed and imaged by using fluorescence microscopy (Carl-Zeiss, Oberkochen, Germany). The intensity of fluorescence staining was measured using ImageJ software (NIH, Bethesda, MD).

#### Fabrication and characterization of 3D hydrogels

GelMA hydrogel was fabricated as previously reported (*108*). Briefly, 10% (w/v) GelMA (Jurassic, Haining, China) solution and a photo-crosslinker, LAP (40mg/ml, Jurassic) were mixed in a ratio of 40:1 at 37 ℃. Then, SF or SS solution was added into hydrogel precursor solution for a final concentration of 0.2% (w/v). A UV irradiation (Wave Length: 365-370nm; Light Strength: 50 mW/cm^2^) for 20 sec was conducted to achieve the photocrosslinking of GelMA hydrogel. The surface morphology of hydrogels was observed and imaged by using a Zeiss EVO 18 Scanning electron microscope (SEM, Carl-Zeiss, Oberkochen, Germany). The pore size of hydrogels in each group was determined by measuring the randomly selected 50 pores from SEM images in ImageJ software (NIH, Bethesda, MD). The porosity of hydrogels was evaluated by using the liquid displacement method as previously reported (*109*). Rheological properties of hydrogels, including storage and loss modulus, was conducted from 0.1 to 100 rad/s at 1% strain by using an MCR 302 rheometer (Anton Paar, Graz, Austria).

#### Mass spectrometry imaging (MSI)

MSI was performed to investigate the metabolites of MSCs in 3D culture. The samples were cut into 10 µm sections at -22℃ using a cryostat microtome (Leica, Wetzlar, Germany) and mounted to conductive indium tin oxide-coated glass slides. After drying in a vacuum for 30 min, slides were stored at -80℃ until MSI analysis. After drying in a vacuum for 30 min, the tissue sections were covered with HCCA (4-Hydroxy-α-cyanocinnamic acid) by Bruker ImagePrep device (Bruker Daltonics, Billerica, MA). MSI was performed using a UltrafleXtreme MALDI TOF MS/MS (Bruker Daltonics) instrument in positive ion mode. The analyzer was operated in reflection mode, and the laser was fired at a repetition rate of 2000 Hz. MSI spatial resolution was set to 100 µm for tissue sections, and each pixel consists of 200 laser shots. The mass spectra data were acquired over the range of m/z 50-1500. The raw tissue MSI data were viewed and processed using Fleximaging 5.0 (Bruker Daltonics) and SCiLS Lab version 2019c (Bruker Daltonics).

#### Rat full-thickness skin defects model

Full-thickness wounds were created in male Sprague Dawley rats (200∼220g) according to the previous report (*110*). The animal experiment protocol was approved by the Animal Experimental Ethical Inspection Committee of Southeast University (20211101001). Briefly, SD rats (n=4 per group) were anesthetized and the skin was cleaned with 75% ethanol after the removal of hair. Full-thickness skin wounds were created by using a biopsy pouch with a diameter of 10 mm on the dorsal of each rat. 0.2 ml GelMA hydrogel incorporated with/without 0.2% (w/v) SF or SS was formed on the local sites of skin wounds after UV radiation for 20 sec (Wave Length: 365-370 nm; Light Strength: 50 mW/cm^2^). For MSC-seeded hydrogels, 2×10^5^ MSCs were encapsulated inside before gelation. To prevent the immune response of human MSCs transplantation in rat model of skin defect, primary rat MSCs (Cyagen Biosciences, Suzhou, China) from bone marrow were used for the in vivo study. Digital images of the skin wounds in SD rats were taken on days 0, 3, 7, 10, and 14, and the wound healing rate of wounds was determined by ImageJ software (NIH, Bethesda, MD). On day 14 post-surgery, all SD rats were sacrificed, and each wound tissue was collected and subsequently cut in half for histological evaluation and proteomic analysis.

#### Histological and immunohistochemical (IHC) staining

The wound tissues from SD rats were fixed in 4% PFA for 24 h, embedded in paraffin, and sectioned into 7 μm thick sections. After deparaffinization and rehydration, the sections were stained with hematoxylin and eosin (H&E) and Masson’s Trichrome (MT) dye. Quantitative analyses for skin thickness, vessel number and area, and hair follicle number were performed from H&E stained images. For IHC staining, the sections were incubated in the first antibody overnight for rabbit anti-COL1 (Proteintech, Wuhan, China), anti-CD31 (Servicebio, Wuhan, China), anti-IL-1β (Servicebio, Wuhan, China), anti-CD206 (Proteintech, Wuhan, China) followed deparaffinization and antigen retrieval, as well as blocking. Subsequently, the sections were stained with HRP conjugated secondary antibody at room temperature for 2 h (Servicebio, Wuhan, China). The DAB substrate system (Servicebio, Wuhan, China) was used for color development. Hematoxylin was applied to the staining of nuclear.

#### Proteomic analysis for protein expression in the wound microenvironments

The wound tissues were collected from SD rats on day 3 post-surgery (n = 3/group). Total proteins were extracted from the collected wound tissue of each group. Subsequently, the LFQ-based proteomic analysis was conducted to investigate the protein expression of the wound environment after SF or SS treatment. The proteins with a P value less than 0.05 after paired comparison were used for GO and KEGG enrichment analyses on the DAVID website. The protein interaction network from DEPs was visualized on the STRING, Cytoscape, and CytoHubba. Other bioinformatic data were analyzed on the OmicStudio Tools (https://www.omicstudio.cn/). The mass spectrometry proteomics data have been deposited to the ProteomeXchange Consortium (http://proteomecentral.proteomexchange.org) via the iProX partner repository with the dataset identifier PXD035603.

#### Statistics

Data were presented as mean ± standard deviation (SD). Student’s t-test, one-way ANOVA, and two-way ANOVA with Tukey’s multiple comparisons were used for statistical analysis, with statistical significance at P < 0.05.

## Acknowledgments

The authors would like to thank Prof. Jie Chao (Southeast University) and Prof. Lixin Wang (Southeast University) for generously providing the cell lines.

## Author contributions

Conceptualization: Wei Zhang, Jianlin Chen, Hongmei Wang

Methodology: Wei Zhang, Yanan Zhang, Jianlin Chen, Hongmei Wang

Investigation: Yanan Zhang, Yue Zhu, Zhicheng Cao, Xinxi Zhao, Chuanquan Liu, Zhixuan Chen, Po Zhang, Baian Kuang, Haotian Zheng

Visualization: Renwang Sheng, Wei Zhang, Yanan Zhang

Supervision: Wei Zhang, Zhimei Wang, Qingqiang Yao,

Writing—original draft: Renwang Sheng, Yanan Zhang, Wei Zhang

Writing—review & editing: Wei Zhang, Renwang Sheng, Jialin Chen, Hongmei Wang

## Data and materials availability

All data that are required to demonstrate the conclusions are showed in the paper and the Supplementary Materials. Additional data in relation to this paper may be requested from the authors.

## References

1. G. M. Abouna, Organ shortage crisis: problems and possible solutions. Transplant Proc 40, 34–38 (2008).

2. R. Langer, J. P. Vacanti, Tissue engineering. Science 260, 920–926 (1993).

3. F. J. O’Brien, Biomaterials & scaffolds for tissue engineering. Materials Today 14, 88–95 (2011).

4. A. Ho-Shui-Ling et al., Bone regeneration strategies: Engineered scaffolds, bioactive molecules and stem cells current stage and future perspectives. Biomaterials 180, 143–162 (2018).

5. Y. Xu et al., Combined chemical and structural signals of biomaterials synergistically activate cell-cell communications for improving tissue regeneration. Acta Biomater 55, 249–261 (2017).

6. C. Holland, K. Numata, J. Rnjak-Kovacina, F. P. Seib, The Biomedical Use of Silk: Past, Present, Future. Adv Healthc Mater 8, e1800465 (2019).

7. B. Kundu, R. Rajkhowa, S. C. Kundu, X. Wang, Silk fibroin biomaterials for tissue regenerations. Adv Drug Deliv Rev 65, 457–470 (2013).

8. Y. Q. Zhang, Applications of natural silk protein sericin in biomaterials. Biotechnol Adv 20, 91–100 (2002).

9. W. Zhang et al., An all-silk-derived functional nanosphere matrix for sequential biomolecule delivery and in situ osteochondral regeneration. Bioact Mater 5, 832–843 (2020).

10. W. Zhang et al., Silk Fibroin Biomaterial Shows Safe and Effective Wound Healing in Animal Models and a Randomized Controlled Clinical Trial. Adv Healthc Mater 6, 1700121 (2017).

11. W. Zhang et al., Sustained Release of TPCA-1 from Silk Fibroin Hydrogels Preserves Keratocyte Phenotype and Promotes Corneal Regeneration by Inhibiting Interleukin-1beta Signaling. Adv Healthc Mater 9, 2000591 (2020).

12. L. Zhang et al., Systematic Review of Silk Scaffolds in Musculoskeletal Tissue Engineering Applications in the Recent Decade. ACS Biomater Sci Eng, (2021).

13. G. H. Altman et al., Silk-based biomaterials. Biomaterials 24, 401–416 (2003).

14. Z. Jiao et al., In Vivo Characterizations of the Immune Properties of Sericin: An Ancient Material with Emerging Value in Biomedical Applications. Macromol Biosci 17, (2017).

15. Y. Song et al., An injectable silk sericin hydrogel promotes cardiac functional recovery after ischemic myocardial infarction. Acta Biomater 41, 210–223 (2016).

16. L. Lamboni, M. Gauthier, G. Yang, Q. Wang, Silk sericin: A versatile material for tissue engineering and drug delivery. Biotechnol Adv 33, 1855–1867 (2015).

17. Y. R. Park et al., NF-kappaB signaling is key in the wound healing processes of silk fibroin. Acta Biomater 67, 183–195 (2018).

18. C. Martinez-Mora et al., Fibroin and sericin from Bombyx mori silk stimulate cell migration through upregulation and phosphorylation of c-Jun. J Tissue Eng Regen M 6, 177–177 (2012).

19. Y. Q. Wang et al., Effect of Electrospun Silk Fibroin-Silk Sericin Films on Macrophage Polarization and Vascularization. Acs Biomater Sci Eng 6, 3502–3512 (2020).

20. N. Bhardwaj, S. C. Kundu, Chondrogenic differentiation of rat MSCs on porous scaffolds of silk fibroin/chitosan blends. Biomaterials 33, 2848–2857 (2012).

21. G. J. Lai, K. T. Shalumon, S. H. Chen, J. P. Chen, Composite chitosan/silk fibroin nanofibers for modulation of osteogenic differentiation and proliferation of human mesenchymal stem cells. Carbohyd Polym 111, 288–297 (2014).

22. H. Y. Han et al., Silk Biomaterials with Vascularization Capacity. Adv Funct Mater 26, 421–432 (2016).

23. S. Bai et al., Silk scaffolds with tunable mechanical capability for cell differentiation. Acta Biomater 20, 22–31 (2015).

24. S. Bhowmick, D. Scharnweber, V. Koul, Co-cultivation of keratinocyte-human mesenchymal stem cell (hMSC) on sericin loaded electrospun nanofibrous composite scaffold (cationic gelatin/hyaluronan/chondroitin sulfate) stimulates epithelial differentiation in hMSCs: In vitro study. Biomaterials 88, 83–96 (2016).

25. G. Griffanti, W. G. Jiang, S. N. Nazhat, Bioinspired mineralization of a functionalized injectable dense collagen hydrogel through silk sericin incorporation. Biomater Sci-Uk 7, 1064–1077 (2019).

26. C. Martinez-Mora et al., Fibroin and Sericin from Bombyx mori Silk Stimulate Cell Migration through Upregulation and Phosphorylation of c-Jun. Plos One 7, (2012).

27. J. K. Carrow et al., Widespread changes in transcriptome profile of human mesenchymal stem cells induced by two-dimensional nanosilicates. P Natl Acad Sci USA 115, E3905–E3913 (2018).

28. Z. Wang, M. Gerstein, M. Snyder, RNA-Seq: a revolutionary tool for transcriptomics. Nat Rev Genet 10, 57–63 (2009).

29. J. K. Carrow et al., Widespread changes in transcriptome profile of human mesenchymal stem cells induced by two-dimensional nanosilicates. Proc Natl Acad Sci U S A 115, E3905–E3913 (2018).

30. P. A. Guerette et al., Accelerating the design of biomimetic materials by integrating RNA-seq with proteomics and materials science. Nat Biotechnol 31, 908–915 (2013).

31. X. Wang, Q. Liu, B. Zhang, Leveraging the complementary nature of RNA-Seq and shotgun proteomics data. Proteomics 14, 2676–2687 (2014).

32. J. K. Carrow et al., Photothermal modulation of human stem cells using light-responsive 2D nanomaterials. Proc Natl Acad Sci U S A 117, 13329–13338 (2020).

33. Z. Othman et al., Comparative proteomic analysis of human mesenchymal stromal cell behavior on calcium phosphate ceramics with different osteoinductive potential. Mater Today Bio 7, 100066 (2020).

34. V. Bunpetch et al., Silicate-based bioceramic scaffolds for dual-lineage regeneration of osteochondral defect. Biomaterials 192, 323–333 (2019).

35. R. Sheng et al., Nanosilicate-Reinforced Silk Fibroin Hydrogel for Endogenous Regeneration of Both Cartilage and Subchondral Bone. Adv Healthc Mater, e2200602 (2022).

36. J. H. Lee, D. W. Song, Y. H. Park, I. C. Um, Effect of residual sericin on the structural characteristics and properties of regenerated silk films. Int J Biol Macromol 89, 273–278 (2016).

37. X. M. Zhang, P. Wyeth, Using FTIR spectroscopy to detect sericin on historic silk. Sci China Chem 53, 626–631 (2010).

38. W. Z. Sun, D. A. Gregory, M. A. Tomeh, X. B. Zhao, Silk Fibroin as a Functional Biomaterial for Tissue Engineering. International Journal of Molecular Sciences 22, (2021).

39. E. Bari, S. Perteghella, S. Farago, M. L. Torre, Association of silk sericin and platelet lysate: Premises for the formulation of wound healing active medications. Int J Biol Macromol 119, 37–47 (2018).

40. A. J. Meinel et al., Optimization strategies for electrospun silk fibroin tissue engineering scaffolds. Biomaterials 30, 3058–3067 (2009).

41. M. Floren et al., Human mesenchymal stem cells cultured on silk hydrogels with variable stiffness and growth factor differentiate into mature smooth muscle cell phenotype. Acta Biomater 31, 156–166 (2016).

42. W. C. Tsai et al., Platelet rich plasma promotes skeletal muscle cell migration in association with up-regulation of FAK, paxillin, and F-Actin formation. Journal of Orthopaedic Research 35, 2506–2512 (2017).

43. J. C. Porter, M. Bracke, A. Smith, D. Davies, N. Hogg, Signaling through integrin LFA-1 leads to filamentous actin polymerization and remodeling, resulting in enhanced T cell adhesion. J Immunol 168, 6330–6335 (2002).

44. E. Bari, S. Perteghella, S. Farago, M. L. Torre, Association of silk sericin and platelet lysate: Premises for the formulation of wound healing active medications. Int J Biol Macromol 119, 37–47 (2018).

45. Y. F. Yan et al., Enhanced Osteogenesis of Bone Marrow-Derived Mesenchymal Stem Cells by a Functionalized Silk Fibroin Hydrogel for Bone Defect Repair. Adv Healthc Mater 8, (2019).

46. X. Hu et al., The influence of elasticity and surface roughness on myogenic and osteogenic-differentiation of cells on silk-elastin biomaterials. Biomaterials 32, 8979–8989 (2011).

47. D. W. Li et al., Silk fibroin/chitosan thin film promotes osteogenic and adipogenic differentiation of rat bone marrow-derived mesenchymal stem cells. J Biomater Appl 32, 1164–1173 (2018).

48. M. Orecchioni et al., Molecular and Genomic Impact of Large and Small Lateral Dimension Graphene Oxide Sheets on Human Immune Cells from Healthy Donors. Adv Healthc Mater 5, 276–287 (2016).

49. Y. Z. Zhu et al., Regulation of macrophage polarization through surface topography design to facilitate implant-to-bone osteointegration. Sci Adv 7, (2021).

50. T. H. Qazi, D. J. Mooney, G. N. Duda, S. Geissler, Biomaterials that promote cell-cell interactions enhance the paracrine function of MSCs. Biomaterials 140, 103–114 (2017).

51. N. Su et al., Fibrous scaffolds potentiate the paracrine function of mesenchymal stem cells: A new dimension in cell-material interaction. Biomaterials 141, 74–85 (2017).

52. R. Y. Huang et al., The topography of fibrous scaffolds modulates the paracrine function of Ad-MSCs in the regeneration of skin tissues. Biomater Sci-Uk 7, 4248–4259 (2019).

53. H. M. Wobma, D. Liu, G. Vunjak-Novakovic, Paracrine Effects of Mesenchymal Stromal Cells Cultured in Three-Dimensional Settings on Tissue Repair. ACS Biomater Sci Eng 4, 1162–1175 (2018).

54. X. Y. Ma et al., Silk Fibroin/Hydroxyapatite Coating Improved Osseointegration of Porous Titanium Implants under Diabetic Conditions via Activation of the PI3K/Akt Signaling Pathway. ACS Biomater Sci Eng 8, 2908–2919 (2022).

55. H. Zhang et al., FN1 promotes chondrocyte differentiation and collagen production via TGF-beta/PI3K/Akt pathway in mice with femoral fracture. Gene 769, 145253 (2021).

56. Q. Han et al., A supramolecular hydrogel based on the combination of YIGSR and RGD enhances mesenchymal stem cells paracrine function via integrin α2β1 and PI3K/AKT signaling pathway for acute kidney injury therapy. Chem Eng J 436, (2022).

57. J. Zhang et al., Magnesium modification of a calcium phosphate cement alters bone marrow stromal cell behavior via an integrin-mediated mechanism. Biomaterials 53, 251–264 (2015).

58. U. Blache, M. M. Stevens, E. Gentleman, Harnessing the secreted extracellular matrix to engineer tissues. Nat Biomed Eng 4, 357–363 (2020).

59. K. H. Vining, D. J. Mooney, Mechanical forces direct stem cell behaviour in development and regeneration. Nat Rev Mol Cell Biol 18, 728–742 (2017).

60. K. Wolf et al., Collagen-based cell migration models in vitro and in vivo. Semin Cell Dev Biol 20, 931–941 (2009).

61. P. Henriet, Z. D. Zhong, P. C. Brooks, K. I. Weinberg, Y. A. DeClerck, Contact with fibrillar collagen inhibits melanoma cell proliferation by up-regulating p27KIP1. Proc Natl Acad Sci U S A 97, 10026–10031 (2000).

62. C. W. Chen et al., Type I and II collagen regulation of chondrogenic differentiation by mesenchymal progenitor cells. J Orthop Res 23, 446–453 (2005).

63. B. Chevallay, D. Herbage, Collagen-based biomaterials as 3D scaffold for cell cultures: applications for tissue engineering and gene therapy. Med Biol Eng Comput 38, 211–218 (2000).

64. C. Piard et al., 3D printed HUVECs/MSCs cocultures impact cellular interactions and angiogenesis depending on cell-cell distance. Biomaterials 222, 119423 (2019).

65. G. Cantarella et al., Nerve growth factor-endothelial cell interaction leads to angiogenesis in vitro and in vivo. Faseb J 16, 1307-+ (2002).

66. L. A. van Meeteren, M. J. Goumans, P. ten Dijke, TGF-beta receptor signaling pathways in angiogenesis; emerging targets for anti-angiogenesis therapy. Curr Pharm Biotechnol 12, 2108–2120 (2011).

67. H. F. Dvorak, Angiogenesis: update 2005. J Thromb Haemost 3, 1835–1842 (2005).

68. C. S. Abhinand, R. Raju, S. J. Soumya, P. S. Arya, P. R. Sudhakaran, VEGF-A/VEGFR2 signaling network in endothelial cells relevant to angiogenesis. J Cell Commun Signal 10, 347–354 (2016).

69. I. Arutyunyan et al., Role of VEGF-A in angiogenesis promoted by umbilical cord-derived mesenchymal stromal/stem cells: in vitro study. Stem Cell Res Ther 7, 46 (2016).

70. W. Katagiri et al., Angiogenesis in newly regenerated bone by secretomes of human mesenchymal stem cells. Maxillofac Plast Reconstr Surg 39, 8 (2017).

71. H. Mayer et al., Vascular endothelial growth factor (VEGF-A) expression in human mesenchymal stem cells: autocrine and paracrine role on osteoblastic and endothelial differentiation. J Cell Biochem 95, 827–839 (2005).

72. S. Lin et al., IGF-1 promotes angiogenesis in endothelial cells/adipose-derived stem cells co-culture system with activation of PI3K/Akt signal pathway. Cell Prolif 50, (2017).

73. C. Y. Ewald, J. N. Landis, J. Porter Abate, C. T. Murphy, T. K. Blackwell, Dauer-independent insulin/IGF-1-signalling implicates collagen remodelling in longevity. Nature 519, 97–101 (2015).

74. P. Hagerty et al., The effect of growth factors on both collagen synthesis and tensile strength of engineered human ligaments. Biomaterials 33, 6355–6361 (2012).

75. G. Grandjean et al., Definition of a Novel Feed-Forward Mechanism for Glycolysis-HIF1alpha Signaling in Hypoxic Tumors Highlights Aldolase A as a Therapeutic Target. Cancer Res 76, 4259–4269 (2016).

76. B. Cruys et al., Glycolytic regulation of cell rearrangement in angiogenesis. Nature Communications 7, (2016).

77. K. De Bock et al., Role of PFKFB3-Driven Glycolysis in Vessel Sprouting. Cell 154, 651–663 (2013).

78. Z. P. Liu et al., Glycolysis links reciprocal activation of myeloid cells and endothelial cells in the retinal angiogenic niche. Sci Transl Med 12, (2020).

79. X. F. Hu, Y. F. Feng, G. Xiang, W. Lei, L. Wang, Lactic acid of PLGA coating promotes angiogenesis on the interface between porous titanium and diabetic bone. J Mater Chem B 6, 2274–2288 (2018).

80. J. Song et al., Lactic Acid Upregulates VEGF Expression in Macrophages and Facilitates Choroidal Neovascularization. Invest Ophthalmol Vis Sci 59, 3747–3754 (2018).

81. M. Rodrigues, N. Kosaric, C. A. Bonham, G. C. Gurtner, Wound Healing: A Cellular Perspective. Physiol Rev 99, 665–706 (2019).

82. P. Bainbridge, Wound healing and the role of fibroblasts. J Wound Care 22, 407–408, 410–412 (2013).

83. Y. Xiao, MiR-486-5p inhibits the hyperproliferation and production of collagen in hypertrophic scar fibroblasts via IGF1/PI3K/AKT pathway. J Dermatolog Treat 32, 973–982 (2021).

84. M. Hetzel, M. Bachem, D. Anders, G. Trischler, M. Faehling, Different effects of growth factors on proliferation and matrix production of normal and fibrotic human lung fibroblasts. Lung 183, 225–237 (2005).

85. J. C. Chen et al., NGF accelerates cutaneous wound healing by promoting the migration of dermal fibroblasts via the PI3K/Akt-Rac1-JNK and ERK pathways. Biomed Res Int 2014, 547187 (2014).

86. Y. R. Yun et al., Fibroblast growth factors: biology, function, and application for tissue regeneration. J Tissue Eng 2010, 218142 (2010).

87. R. Augustine et al., Electrospun polycaprolactone membranes incorporated with ZnO nanoparticles as skin substitutes with enhanced fibroblast proliferation and wound healing. Rsc Adv 4, 24777–24785 (2014).

88. G. Han et al., Nitric Oxide-Releasing Nanoparticles Accelerate Wound Healing by Promoting Fibroblast Migration and Collagen Deposition. Am J Pathol 180, 1465–1473 (2012).

89. Y. F. Lu et al., Engineering Bacteria-Activated Multifunctionalized Hydrogel for Promoting Diabetic Wound Healing. Advanced Functional Materials 31, (2021).

90. S. P. Herbert, D. Y. R. Stainier, Molecular control of endothelial cell behaviour during blood vessel morphogenesis. Nat Rev Mol Cell Bio 12, 551–564 (2011).

91. S. Shigematsu et al., IGF-1 regulates migration and angiogenesis of human endothelial cells. Endocr J 46 Suppl, S59–62 (1999).

92. G. Cantarella et al., Nerve growth factor-endothelial cell interaction leads to angiogenesis in vitro and in vivo. FASEB J 16, 1307–1309 (2002).

93. M. Rodrigues, N. Kosaric, C. A. Bonham, G. C. Gurtner, Wound Healing: A Cellular Perspective. Physiol Rev 99, 665–706 (2019).

94. X. Chen et al., IL-6 induced M1 type macrophage polarization increases radiosensitivity in HPV positive head and neck cancer. Cancer Lett 456, 69–79 (2019).

95. M. Heusinkveld et al., M2 macrophages induced by prostaglandin E2 and IL-6 from cervical carcinoma are switched to activated M1 macrophages by CD4+ Th1 cells. J Immunol 187, 1157–1165 (2011).

96. D. Montoya et al., IL-32 is a molecular marker of a host defense network in human tuberculosis. Sci Transl Med 6, (2014).

97. A. Osman et al., M-CSF Inhibits Anti-HIV-1 Activity of IL-32, but They Enhance M2-like Phenotypes of Macrophages. J Immunol 192, 5083–5089 (2014).

98. M. Orecchioni, Y. Ghosheh, A. B. Pramod, K. Ley, Macrophage Polarization: Different Gene Signatures in M1(LPS+) vs. Classically and M2(LPS-) vs. Alternatively Activated Macrophages. Front Immunol 10, (2019).

99. J. Giri, R. Das, E. Nylen, R. Chinnadurai, J. Galipeau, CCL2 and CXCL12 Derived from Mesenchymal Stromal Cells Cooperatively Polarize IL-10+Tissue Macrophages to Mitigate Gut Injury. Cell Rep 30, 1923-+ (2020).

100. J. Wu et al., TNFSF9 promotes metastasis of pancreatic cancer through Wnt/Snail signaling and M2 polarization of macrophages. Aging-Us 13, 21571–21586 (2021).

101. C. C. Zhao et al., TNFSF15 facilitates differentiation and polarization of macrophages toward M1 phenotype to inhibit tumor growth. Oncoimmunology 11, 2032918 (2022).

102. S. Ansari, S. Pouraghaei Sevari, C. Chen, P. Sarrion, A. Moshaverinia, RGD-Modified Alginate-GelMA Hydrogel Sheet Containing Gingival Mesenchymal Stem Cells: A Unique Platform for Wound Healing and Soft Tissue Regeneration. ACS Biomater Sci Eng 7, 3774–3782 (2021).

103. N. Kasoju, U. Bora, Silk fibroin in tissue engineering. Adv Healthc Mater 1, 393–412 (2012).

104. W. Zhang et al., Silk Fibroin Biomaterial Shows Safe and Effective Wound Healing in Animal Models and a Randomized Controlled Clinical Trial. Adv Healthc Mater 6, (2017).

105. G. Shaikh, B. Cronstein, Signaling pathways involving adenosine A2A and A2B receptors in wound healing and fibrosis. Purinergic Signal 12, 191–197 (2016).

106. D. N. Rockwood et al., Materials fabrication from Bombyx mori silk fibroin. Nat Protoc 6, 1612–1631 (2011).

107. B. B. Mandal, A. S. Priya, S. C. Kundu, Novel silk sericin/gelatin 3-D scaffolds and 2-D films: fabrication and characterization for potential tissue engineering applications. Acta Biomater 5, 3007–3020 (2009).

108. K. Yue et al., Synthesis, properties, and biomedical applications of gelatin methacryloyl (GelMA) hydrogels. Biomaterials 73, 254–271 (2015).

109. D. H. Liang, Z. Lu, H. Yang, J. T. Gao, R. Chen, Novel Asymmetric Wettable AgNPs/Chitosan Wound Dressing: In Vitro and In Vivo Evaluation. Acs Appl Mater Inter 8, 3958–3968 (2016).

110. X. Yao, et al., In-cytoplasm mitochondrial transplantation for mesenchymal stem cells engineering and tissue regeneration. Bioeng Transl Med 7, e10250 (2022).

