## Supplementary material for "Multiomics reveal that silk fibroin and sericin differentially potentiate the paracrine functions of mesenchymal stem cells and enhance tissue regeneration": Materials and methods; Supplemental Figure S1-S17; Supplemental Table 1.

### Supplementary Information

#### Supplementary figures

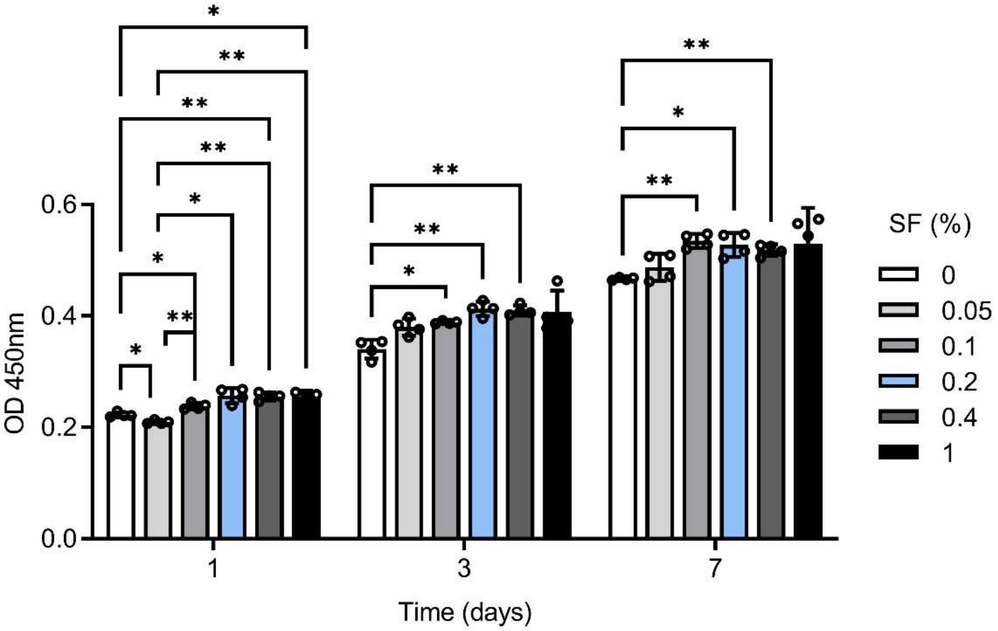


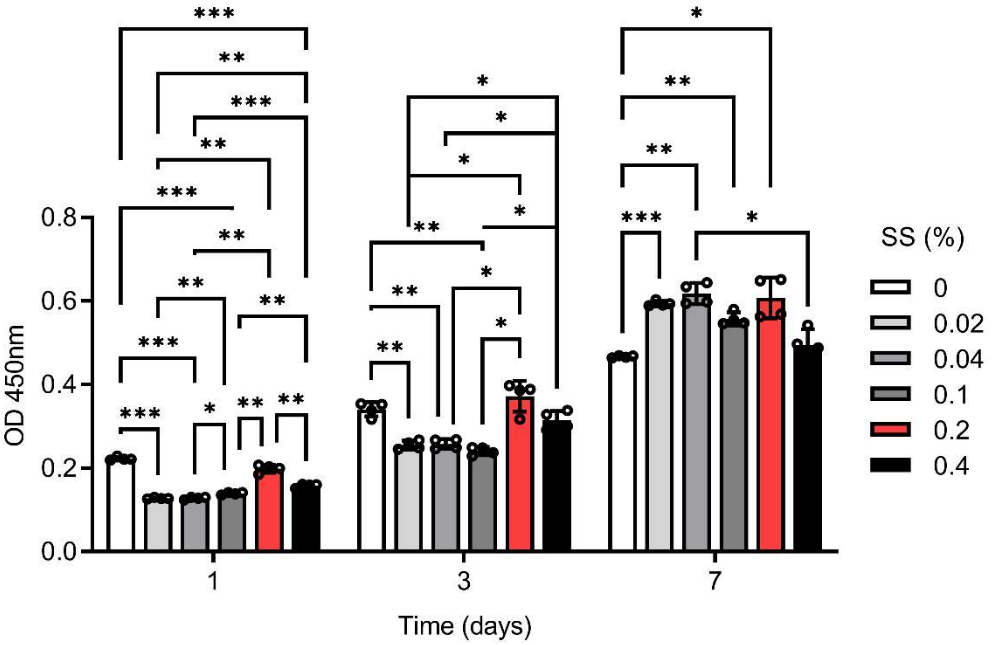


**Figure S1. Cell proliferation of MSCs treated with different concentrations of SF or SS.** The results were presented as mean ± SD. All pairwise comparisons with P values<0.05 are displayed. *P<0.05, **P<0.01, ***P<0.001.

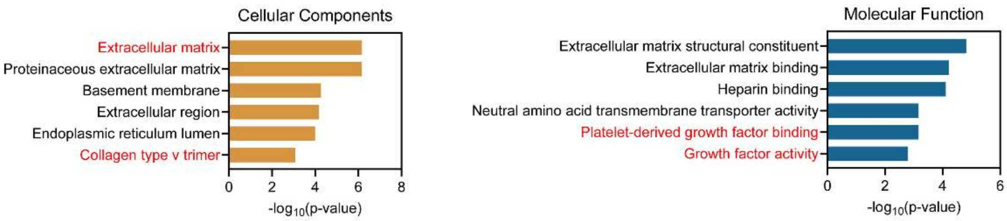


**Figure S2. Key GO terms significantly enriched in CC and MF (SF vs Ctrl) from up-regulated DEGs identified by RNA-seq.**

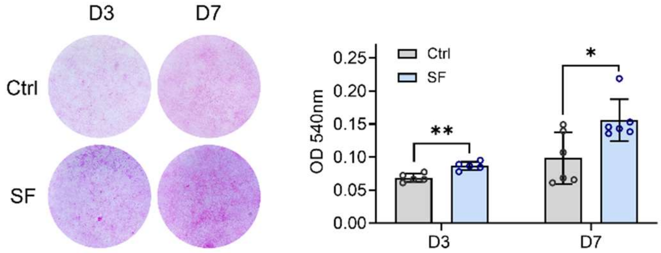


**Figure S3. Representative images and quantitative analysis of picrosirius red staining of MSCs with/without SF treatment.** *P<0.05, **P<0.01.

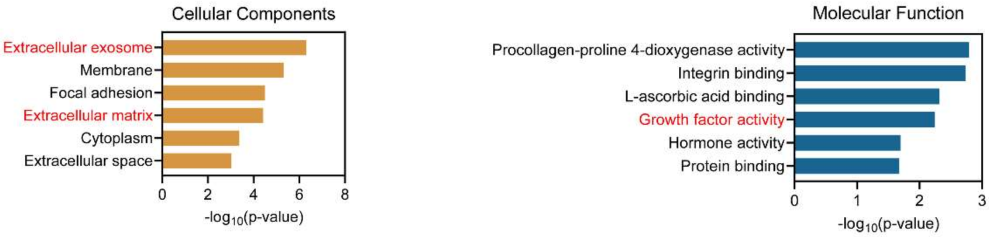

**Figure S4. Key GO terms significantly enriched in CC and MF (SS vs Ctrl) from up-regulated DEGs identified by RNA-seq.**

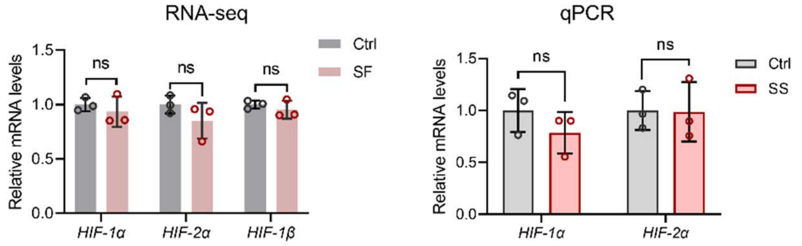


**Figure S5. The expression levels of HIF-1 signaling pathway related genes by RNA-seq and qPCR.**

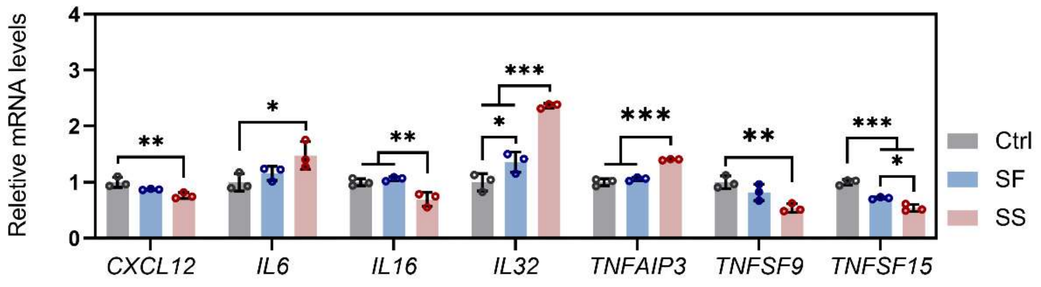


**Figure S6. Gene expression of inflammatory factors in MSCs with different treatments identified by RNA-seq.** The results were presented as mean ± SD. *P<0.05, **P<0.01, ***P<0.001.

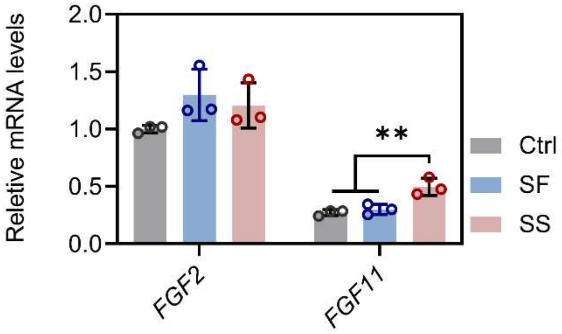


**Figure S7. Gene expression of *FGF2* and *FGF11* in MSCs with different treatments identified by RNA-seq.** The results were presented as mean ± SD. **P<0.01.

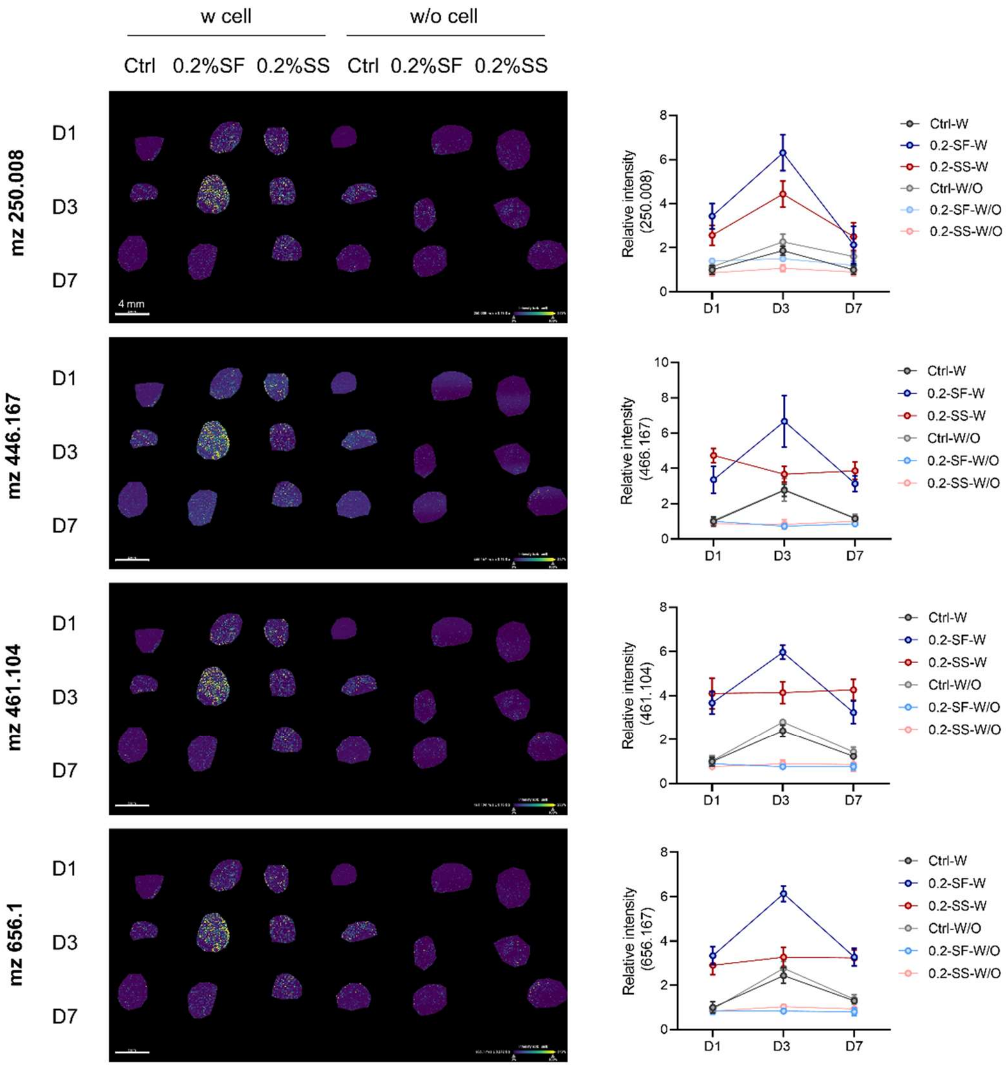


**Figure S8. Spatial distribution and quantitative analysis of cell metabolites of MSCs in 3D hydrogels after different treatments by MSI.** Scale bars = 4 mm.

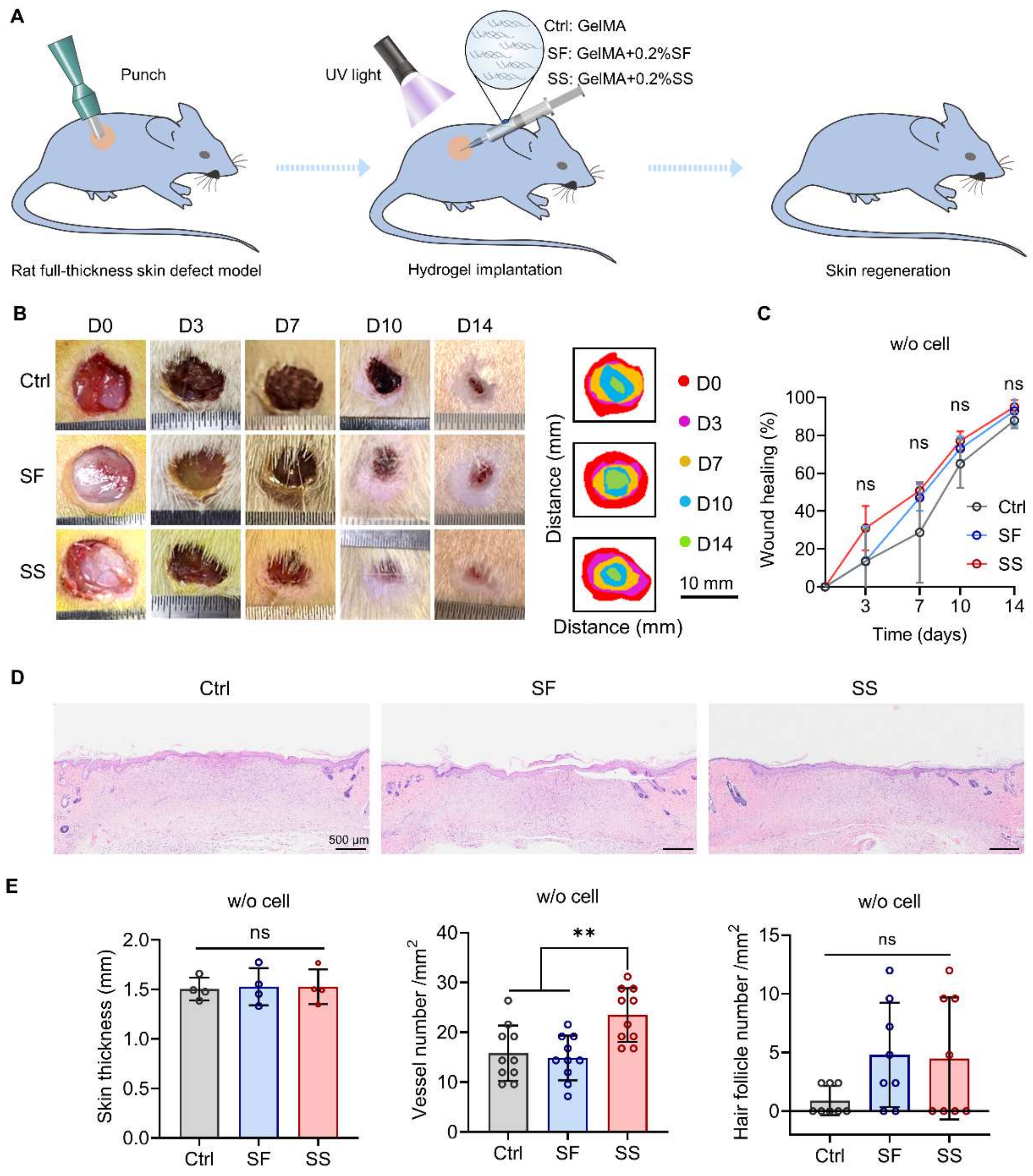


**Figure S9. The effect of SF or SS on skin repair and regeneration in vivo.** (A) Schematic of the animal study procedure. Unseeded GelMA hydrogels incorporated with/without 0.2% (w/v) SF or SS were applied to each wound. (B) Gross morphology of the wound healing process within 14 days. (C) Quantitative analysis of wound healing rate in different groups. (D) H&E staining of regenerated skin tissues in different groups on day 14. Scale bars = 500 μm. (E) Quantitative analysis of skin thickness, vessel number, and hair follicle number from H&E stained images. The results were presented as mean ± SD. **P<0.01.

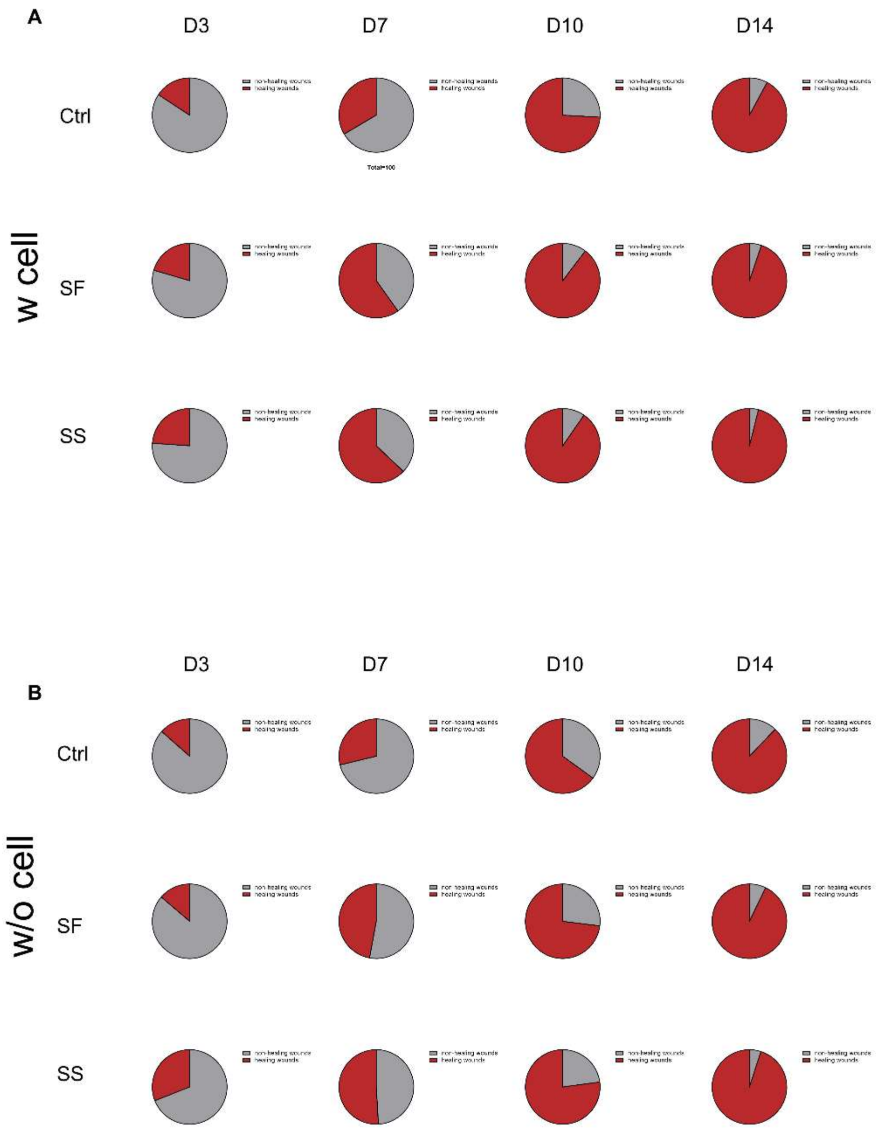


**Figure S10. Fractions of wound healing by different treatments with (A) or without MSCs (B) on days 3, 7, 10, and 14.**

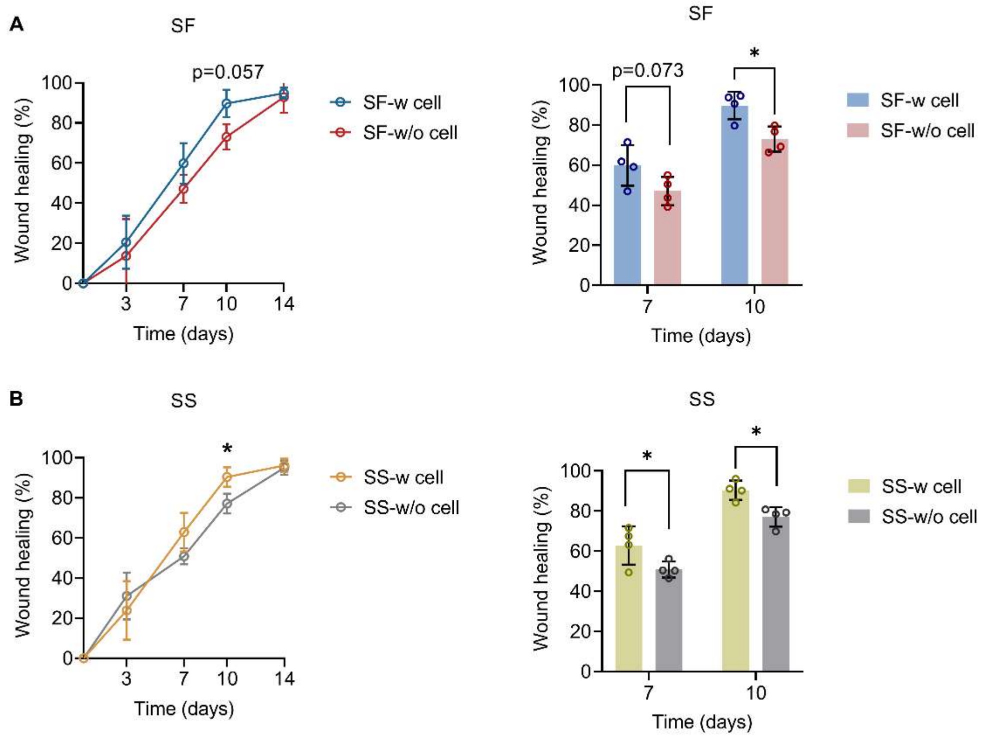


**Figure S11. Comparison of wound healing ratio by SF and SS treatments with/without MSCs.** The results were presented as mean ± SD. *P<0.05.

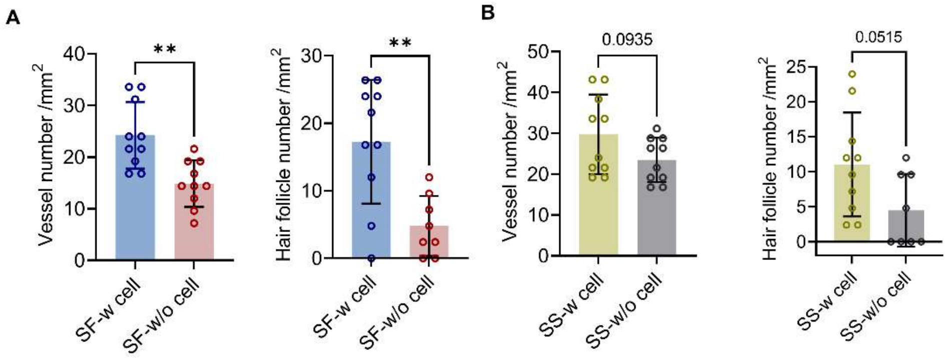


**Figure S12. Comparison of vessel and hair follicle number by SF and SS treatments with/without MSCs.** The results were presented as mean ± SD. **P<0.01.

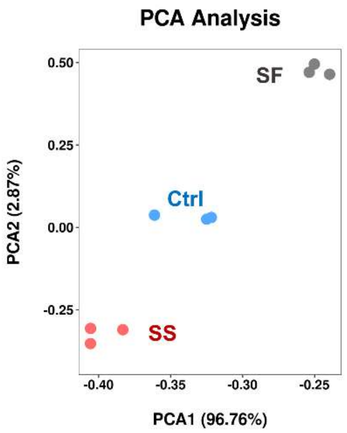


**Figure S13. PCA plot of proteomic data from tissue samples in different groups.**

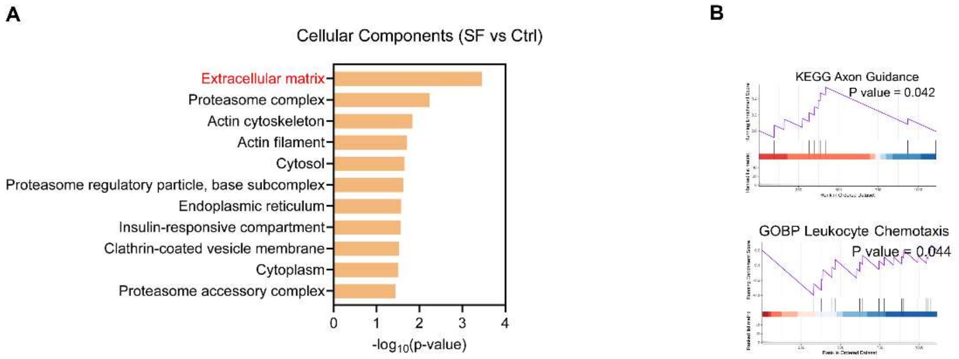


**Figure S14. Functional insights of DEPs identified from tissue samples by proteomic analysis in SF vs Ctrl.** (A) Key GO terms for CC significantly enriched in SF vs Ctrl. (B) GSEA plots of GO terms associated with wound healing (SF vs Ctrl).

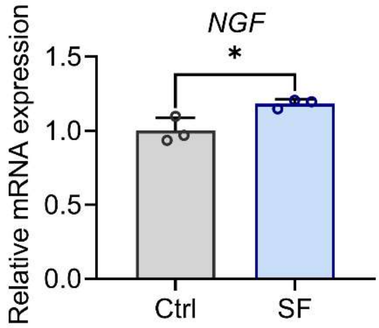


**Figure S15. Gene expression of *NGF* in MSCs after different treatments detected by RNA-seq.** The results were presented as mean ± SD. *P<0.05.

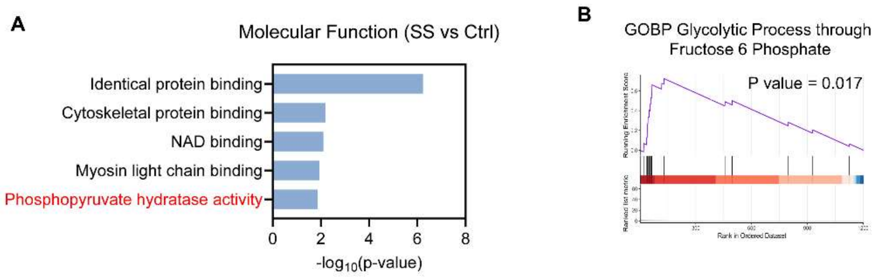


**Figure S16. Functional insights of DEPs identified from tissue samples by proteomic analysis in SS vs Ctrl.** (A) Key GO terms for CC significantly enriched in SS vs Ctrl. (B) GSEA plots of GO terms associated with glycolysis (SS vs Ctrl).

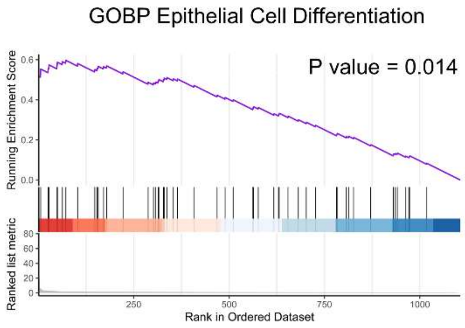


**Figure S17. Functional insights of DEPs identified from tissue samples by proteomic analysis in SF vs SS.**

#### Supplementary table

**Table S1. List of primers for qPCR analysis.**

| Species | Gene | Forward & Reverse | Sequence (5'- 3') | Product length (bp) |
|---|---|---|---|---|
| Human | *β-ACTIN* | Forward | AGCGAGCATCCCCCAAAGTT | 285 |
| | | Reverse | GGGCACGAAGGCTCATCATT | |
| Human | *VEGFA* | Forward | CTTGCCTTGCTGCTCTACC | 201 |
| | | Reverse | CACACAGGATGGCTTGAAG | |
| Human | *IGF-1* | Forward | AGGAAGTACATTTGAAGAACGCAAGT | 104 |
| | | Reverse | CCTGCGGTGGCATGTCA | |
| Human | *NGF* | Forward | GGCAGACCCGCAACATTACT | 135 |
| | | Reverse | CACCACCGACCTCGAAGTC | |

| | | | | |
|---|---|---|---|---|
| Human | *ENO1* | Forward | GTACCGCCACATCGCTGACTTG | 88 |
| | | Reverse | AGCATGAGAACCGCCATTGATGAC | |
| Human | *ENO2* | Forward | GGGAACTCAGACCTCATCCTG | 72 |
| | | Reverse | CTTGTTGCCAGCATGAGAGC | |
| Human | *PDK1* | Forward | ACCAGGACAGCCAATACAAG | 183 |
| | | Reverse | CCTCGGTCACTCATCTTCAC | |
| Human | *HK1* | Forward | GGACTGGACCGTCTGAATGT | 100 |
| | | Reverse | ACAGTTCCTTCACCGTCTGG | |
| Human | *PFKFB3* | Forward | ATCTACCTGAACGTGGAGTCCGTCTG | 270 |
| | | Reverse | TCAGTGTTTCCTGGAGGAGTCAGC | |
| Human | *LDHA* | Forward | TATGGAGTGGAATGAATGTTGC | 238 |
| | | Reverse | CCCTTAATCATGGTGGAAACTC | |
| Human | *SLC2A1* | Forward | CGGGCCAAGAGTGTGCTAAA | 283 |
| | | Reverse | TGACGATACCGGAGCCAATG | |
| Human | *HIF-1α* | Forward | TTCCAGTTACGTTCCTTCGATCA | 76 |
| | | Reverse | TTTGAGGACTTGCGCTTTCA | |
| Human | *HIF-2α* | Forward | AGCAGCTGGAGAGCAAGAAG | 124 |

| | | | | |
|---|---|---|---|---|
| | | Reverse | ATGGAAGAGAGAGGGGTGCT | |
| Mouse | *GAPDH* | Forward | GCAAGTTCAACGGCACAG | 141 |
| | | Reverse | CGCCAGTAGACTCCACGAC | |
| Mouse | *IL-1β* | Forward | AAGGAGAACCAAGCAACGACAAAA | 213 |
| | | Reverse | TGGGGAACTCTGCAGACTCAAACT | |
| Mouse | *TNF-α* | Forward | CGTCAGCCGATTTGCTATCT | 206 |
| | | Reverse | CGGACTCCGCAAAGTCTAAG | |
| Mouse | *iNOS* | Forward | CCTGTGTTCCACCAGGAGAT | 247 |
| | | Reverse | CCCTGGCTAGTGCTTCAGAC | |
| Mouse | *CD206* | Forward | AGCTTCATCTTCGGGCCTTTG | 178 |
| | | Reverse | GGTGACCACTCCTGCTGCTTTAG | |
| Mouse | *ARG-1* | Forward | GTGAAGAACCCACGGTCTGT | 209 |
| | | Reverse | CTGGTTGTCAGGGGAGTGTT | |
| Mouse | *IL-10* | Forward | CCAAGCCTTATCGGAAATGA | 162 |
| | | Reverse | TTTTCACAGGGGAGAAATCG | |
